# PP4 phosphatase cooperates in recombinational DNA repair by enhancing double-strand break end resection

**DOI:** 10.1101/470427

**Authors:** María Teresa Villoria, Pilar Gutiérrez-Escribano, Facundo Ramos, Esmeralda Alonso-Rodríguez, Eva Merino, Alex Montoya, Holger Kramer, Luis Aragón, Andrés Clemente-Blanco

## Abstract

The role of Rad53 in response to a DNA lesion is central for the accurate orchestration of the DNA damage response. Rad53 activation relies on its phosphorylation by the Mec1 kinase and its own auto-phosphorylation in a manner dependent on the adaptor Rad9. While the mechanism behind Rad53 phosphorylation and activation has been well documented, less is known about the processes that counteract its kinase activity during the response to DNA damage. Here, we describe that PP4 phosphatase dephosphorylates Rad53 during the repair of a double-strand break, a process that impacts on the phosphorylation status of multiple factors involved in the DNA damage response. PP4-dependent Rad53 dephosphorylation stimulates DNA end resection by relieving the negative effect that Rad9 exerts over the Sgs1/Dna2 exonuclease complex. Consequently, elimination of PP4 activity affects DNA resection and repair by single-strand annealing, defects that are bypassed by reducing the hyper-phosphorylation state of Rad53 observed in the absence of the phosphatase. These results confirm that Rad53 is one of the main targets of PP4 during the repair of a DNA lesion and demonstrate that the attenuation of its kinase activity during the initial steps of the repair process is essential to efficiently enhance recombinational DNA repair pathways that depend on long-range resection.

## Introduction

Cells are constantly threatened by different sources of DNA damage that can lead to the accumulation of genetic errors that contribute to genome instability. In order to maintain the integrity of the genomic material, cells have developed highly conserved and sophisticated mechanisms that survey their genome, detect DNA lesions and repair them in order to avoid the propagation of mutations. These pathways, collectively known as the DNA damage response (DDR), activate the expression of genes required for DNA repair and trigger checkpoints that inhibit replication and segregation of damaged DNA (Harper and Elledge, 2007). One of the most hazardous types of DNA lesions is the double-strand break (DSB), produced when the two complementary strands of the double helix are severed simultaneously. DSBs are repaired either by direct ligation of the broken ends, as in non-homologous end joining (NHEJ), or by mechanisms that rely on the searching of homologous sequences to regenerate the original DNA molecule, as DSB repair (DSBR), break-induced replication (BIR) or single-strand annealing (SSA) (Symington et al., 2014). Homology-dependent DSB repair is always initiated by the nucleolytic degradation of the 5′-ended strand to generate single-stranded DNA (ssDNA), a process known as resection (Mimitou and Symington, 2009). Resection is activated by two sequential events, an initial processing carried out by the MRX complex (Mre11-Rad50-Xrs2) that together with Sae2 produce a short 3′ overhang ssDNA (Gobbini et al., 2016) followed by a long-range resection triggered by two redundant pathways involving the Sgs1/Dna2-Top3-Rmi1 complex and the Exo1 nuclease (Zhu et al., 2008). Long resection gives rise to extensive ssDNA tracks that are covered by RPA and the DNA damage checkpoint machinery in order to couple the repair process with cell cycle progression (Ciccia and Elledge, 2010). In *Saccharomyces cerevisiae*, the DNA damage checkpoint is mainly controlled by the phosphatidyl inositol kinase-like kinase (PIKK) Mec1, the ortholog of ATR in mammals (Durocher and Jackson, 2001). Upon activation, ssDNA-bound Mec1 channels the checkpoint signal to other kinases such as Rad53 or Chk1 (Chk2 and Chk1 in mammals, respectively). These two kinases are responsible for the phosphorylation of multiple downstream targets required for the execution of different biological processes encompassed in the DDR, including a G2/M arrest which allows cells to accomplish the repair of the lesion before re-entry in the division cycle.

Rad53 is involved in multiple and diverse functions along the DNA damage response as replisome stabilization, expression of repair genes, inhibition of late origin firing, suppression of recombination at stalled forks, fork resumption and repair (Branzei and Foiani, 2006). Activation of Rad53 in response to a DNA lesion comprises a complex mechanism that relies on the conserved checkpoint adaptor Rad9, a protein that shares functional and structural properties with three human mediators BRCA1, MDC1 and 53BP1 (Zhou and Elledge, 2000). After generation of a DNA break Rad9 is associated to damaged chromatin via its BRCT and TUDOR domains, which anchor the protein to Ser129-phosphorylated histone H2A (γH2A) (Hammet et al., 2007). DNA-bound Rad9 is phosphorylated at the DSB vicinity by the cycling-dependent kinase (CDK) (Abreu et al., 2013) and Mec1 (Schwartz et al., 2002) to create a specific binding surface for the Rad53-FHA domain (Sun et al., 1998; Sweeney et al., 2005). Recruitment of Rad53 to Rad9 stimulates its phosphorylation by Mec1 to become pre-activated. Besides, Rad53 like other kinases, has the ability to phosphorylate itself. In this regard, it has been proposed that the accumulation of Rad53 at the break site enhances its auto-phosphorylation and consequently, its full activation (Pellicioli and Foiani, 2005). Once activated, Rad53 is liberated from the Rad9 complex resulting in the amplification of the checkpoint signal throughout the cell (Gilbert et al., 2001).

Even though Rad53 phosphorylation and activation have been extensively studied in the last years, less is known about the mechanisms controlling Rad53 dephosphorylation and inhibition. Interestingly, appearance of unphosphorylated Rad53 after checkpoint inactivation does not require new protein synthesis, suggesting that the activity of the kinase is regulated by dephosphorylation (Tercero et al., 2003). Up to date, a few number of phosphatases have been involved in Rad53 dephosphorylation. Both PP1 and PP2C were described as protein phosphatases necessary to enhance checkpoint inactivation and cell cycle re-entry by targeting Rad53. However, while PP1 is needed during the recovery from HU treatment (Bazzi et al., 2010), PP2C is required for cell cycle re-entry upon induction of a DSB (Leroy et al., 2003), suggesting that different phosphatases counteract Rad53 phosphorylation depending on the type of lesion generated. Recently, new data coming from different organisms have demonstrated that protein phosphatase 4 (PP4) is also a key regulator of the DNA damage response. PP4 is an ubiquitous serine/threonine phosphatase that regulates many cellular functions (Cohen et al., 2005). It is composed by the catalytic subunit Pph3 (Ppp4c in mammals) and several regulatory subunits that control the specificity of the holoenzyme. Similarly to PP1 and PP2C, PP4 has the ability to dephosphorylate Rad53, in this case after MMS treatment (O’Neill et al., 2007). Interestingly, the role of PP4 in the DNA damage response is not only limited to modulate Rad53 phosphorylation. Dephosphorylation of H2A by PP4 in both yeast and human cells has been linked to an efficient recovery from the DNA damage checkpoint arrest (Chowdhury et al., 2008; Keogh et al., 2006; Nakada et al., 2008). Supporting these observations PP4 was found to regulate many Mec1 targets in response to HU treatment, suggesting that this phosphatase is important to balance the phosphorylation state of multiple substrates in response to replication stress (Hustedt et al., 2015). In mammalian cells, PP4 has been implicated in the binding of 53BP1 to chromatin during the formation of a DNA lesion (Lee et al., 2014) and in promoting cell cycle recovery from the G2/M arrest by dephosphorylating KAP1 (Lee et al., 2012). Remarkably, the function of PP4 in the DDR is not only restricted to enhance checkpoint inactivation and cell cycle re-entry, but also to repair. In this regard it has been shown that in the budding yeast, PP4 collaborates with PP2C to stimulate DSB repair by homologous recombination (Kim et al., 2011). Moreover, experiments in human cells have demonstrated that dephosphorylation of RPA by PP4 facilitates recombinational DNA repair (Lee et al., 2010). Overall, these results indicate that PP4 has the ability to control multiple aspects of the DNA damage response, including DNA repair, DNA damage checkpoint and cell cycle recovery.

Here, we describe that PP4 is involved in the recombinational repair of DSBs by modulating the steady-state phosphorylation of multiple factors involved in the DNA damage response. This is attained by the ability of the phosphatase to counteract Rad53 phosphorylation during the repair of a DNA lesion. Rad53 attenuation enhances DNA end resection in a process largely dependent on the Sgs1/Dna2 pathway. Importantly, PP4-dependent resection is crucial for the correct execution of recombinational pathways that rely specifically on the efficiency of the resection process. Accordingly, lack of PP4 activity causes defects in DNA end resection and repair by SSA. Interestingly, both defects are suppressed by reducing the hyper-phosphorylation levels of Rad53, demonstrating that Rad53 is the major effector of the phosphatase during the repair of a DNA break. All together, we propose that Rad53 inhibition by PP4 is not only restricted to silence the DDR once the DNA lesion has been fixed, but also during the initial steps of the repair process in order to accurate execute recombinational repair pathways that rely on long-range resection for its accomplishment.

## Results

### PP4 facilitates DNA repair by enhancing SSA/BIR

As a first approach to determine the role of PP4 in the DNA damage response in *S. cerevisiae*, we verified the effect of different DNA-damaging agents on the growth of cells lacking the catalytic subunit Pph3. Disruption of the *PPH3* gene gave rise to an increase in sensitivity to several genotoxic drugs, including the UV-mimic 4-nitroquinoline-1-oxide (4-NQO), the alkylating agent methyl methanesulphonate (MMS) and the radiomimetic drug phleomycin (Sup. Fig. 1A,B). The high lethality observed in *pph3*Δ cells exposed to all the DNA-damaging agents tested advises for a general role of PP4 in controlling the DNA damage checkpoint and/or DNA repair mechanisms. Interestingly, *pph3*Δ cells were not sensitive to the ribonucleotide reductase inhibitor hydroxyurea (HU), indicating that PP4 activity is not required to maintain cell viability in response to replicative stress (Sup. Fig. 1A,B). Even though PP4 was originally described for its role in DNA damage checkpoint deactivation and cell cycle re-entry (Keogh et al., 2006; Nakada et al., 2008; O’Neill et al., 2007; Szyjka et al., 2008), there are emerging evidences involving the phosphatase in the repair of DNA lesions (Chowdhury et al., 2008; Kim et al., 2011; Lee et al., 2010). To gain insights into a possible role of PP4 in the repair of DSBs, we first monitored the efficiency of the SSA/BIR repair pathways in the presence or absence of its catalytic subunit Pph3 by using the YMV80 background. This strain contains a 117-base-pair HO-endonuclease recognition sequence within the *LEU2* locus on chromosome III and a 3′ fragment of the *LEU2* gene with homology only to the right end of the break (*U2*) placed 25 Kb downstream the HO cut site (Fig. 1A) (Jain et al., 2009). It has been previously shown that this HO-induced DSB is repaired by SSA (Jain et al., 2009) in a Rad51-independent manner, although it can also be repaired by Rad51-dependent intra-chromosomal BIR (Fig. 1A). The HO endonuclease, regulated by the control of a galactose inducible promoter, was expressed in asynchronous cell cultures and samples were taken up to 24 hours to determine the efficiency of the repair by Southern blot. Elimination of *PPH3* resulted in a delay in the formation of the repair product compared to wild-type cells (Fig. 1B). FACS analysis showed that while both wild-type and *pph3*Δ cells timely execute the G2/M arrest due to the DNA damage checkpoint activation, disruption of *PPH3* resulted in a notorious delay in cell cycle re-entry, correlating with their incapacity to timely repair the DNA lesion (Fig. 1B). Supporting this observation, a *pph3*Δ mutant displayed a higher sensitivity than the wild-type strain when platted on galactose-containing media (Sup. Fig. 1C). These experiments demonstrate that PP4 is necessary for the correct execution of SSA/BIR during the repair of a single induced DSB.

**Figure 1.**
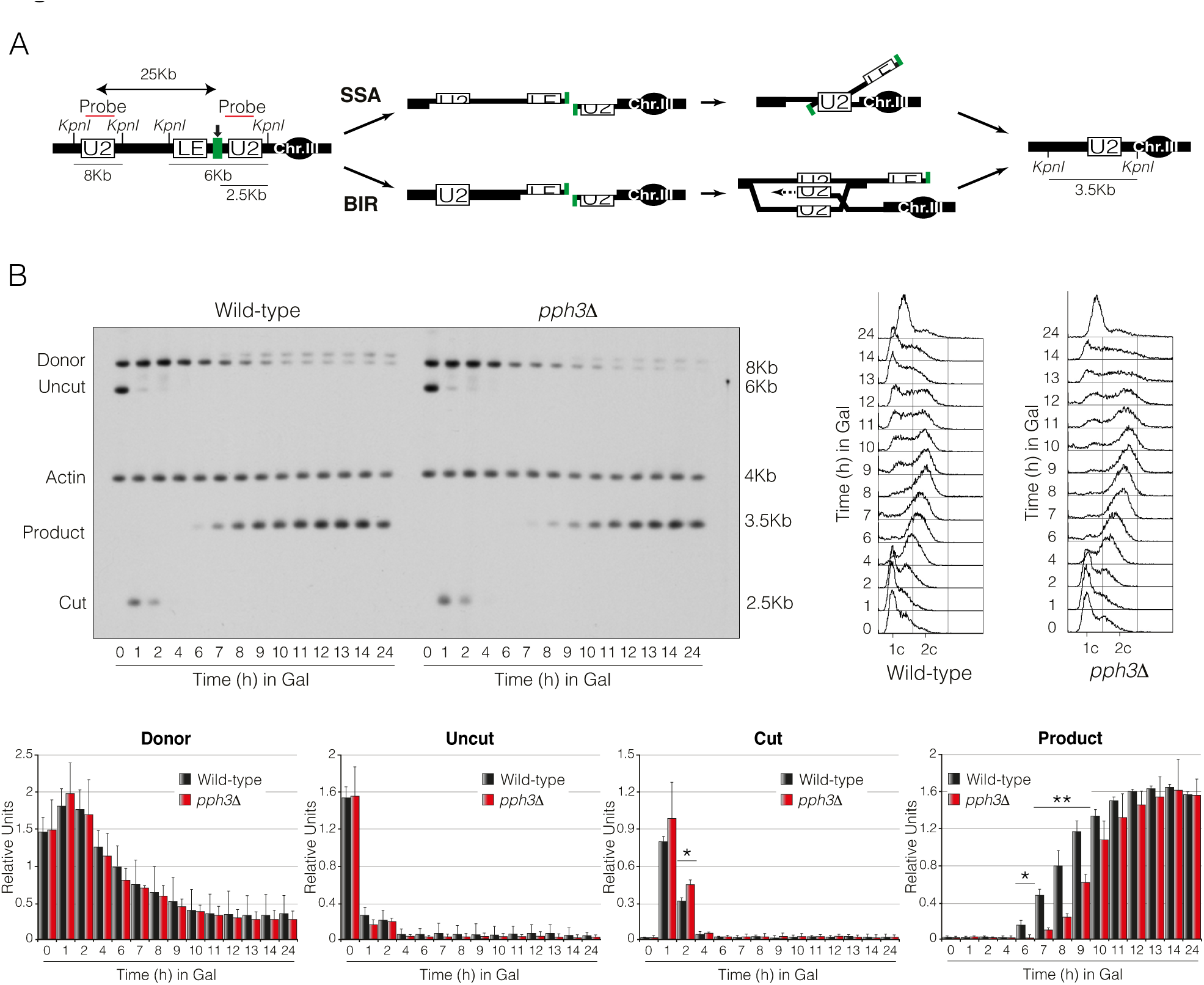
Pph3 is required for DNA repair by homologous recombination. A) Schematic representation depicting relevant genomic structure of the strain used to assess DNA repair by SSA/BIR. The location of an *U2* probe and the restriction endonuclease cleavage sites used for Southern blot analysis to detect repair product formation are shown. Arrow indicates the localization of the double-strand break. B) Physical analysis of wild-type and *pph3Δ* mutant strains carrying the DNA repair system depicted in A). Cells were grown overnight in YP-Raffinose before adding galactose to induce HO expression, thus producing a DSB at the *LEU2* locus on chromosome III. Samples were taken at different time points and genomic DNA was extracted, digested with *Kpn*I and analysed by Southern blot. Blots were hybridized with an *U2* probe DNA sequence and an *ACT1* probe was used as a loading control. FACS profiles for both strains are shown. Graphs show the mean ± SD of the band signals from three independent experiments. All data were normalized using actin as a loading control. Replicates were averaged and statistical significance of differences assessed by a two-tailed unpaired Student’s t-test.

### Identification of PP4 targets in the DNA damage response

In order to identify potential PP4 targets responsible for its function in DNA repair, we performed quantitative phosphoproteomic by mass spectrometry (MS) of samples taken from the experiment described above. Precisely, we screened for variations in the levels of peptide phosphorylation between a wild-type strain and a *pph3Δ* mutant at time 0, 6 and 12 hours after expressing the HO endonuclease. As previously shown in figure 1B, six hours after expressing the HO endonuclease there is still no repaired product formation. Under this condition, those proteins containing increased phosphorylation levels in the absence of Pph3 will reflect a putative target of the phosphatase directly involved in DNA repair regulation. Twelve hours after the induction of the DSB, both wild-type and *pph3Δ* cells have already repaired the DNA lesion, rendering PP4-dependent phosphorylation changes to a function exclusively associated to cell cycle re-entry. From the 11,974 phosphopeptides detected, we isolated those whose phosphorylation was enriched in the absence of PP4 activity and their corresponded proteins were grouped into different GO categories depending on their molecular function (Fig. 2A). A total of 180 proteins with a differential phosphorylation state between the wild-type and *pph3Δ* mutant fell into the DNA damage response category. Within this group, quantitative comparison between the wild-type and the *pph3Δ* strain was applied to classify the discovered targets depending on their averaged phosphorylation across the experimental conditions. A summary containing the most relevant DNA damage proteins identified is depicted in figure 2B. Among the potential PP4 substrates isolated were proteins previously characterized as bona-fide targets of the phosphatase, such as RPA (Lee et al., 2010), histone H2A (Chowdhury et al., 2008; Keogh et al., 2006) or Rad53 (O’Neill et al., 2007; Szyjka et al., 2008). It is important to remark that lack of PP4 activity increases phosphorylation of most of these targets by 6 hours from the DSB induction, when there is still no accumulation of the repair product (Fig. 1B). This suggests that PP4 activity might be required to counterbalance the effect of DDR kinases during the repair of the DNA lesion. Still, we also detected high levels of protein phosphorylation of several DNA damage response proteins in the absence of PP4 activity by 12 hours after the induction of the HO break, a time-point where cells have already accomplished the repair process (Fig. 1B), reflecting its role in checkpoint deactivation and cell cycle re-entry.

**Figure 2.**
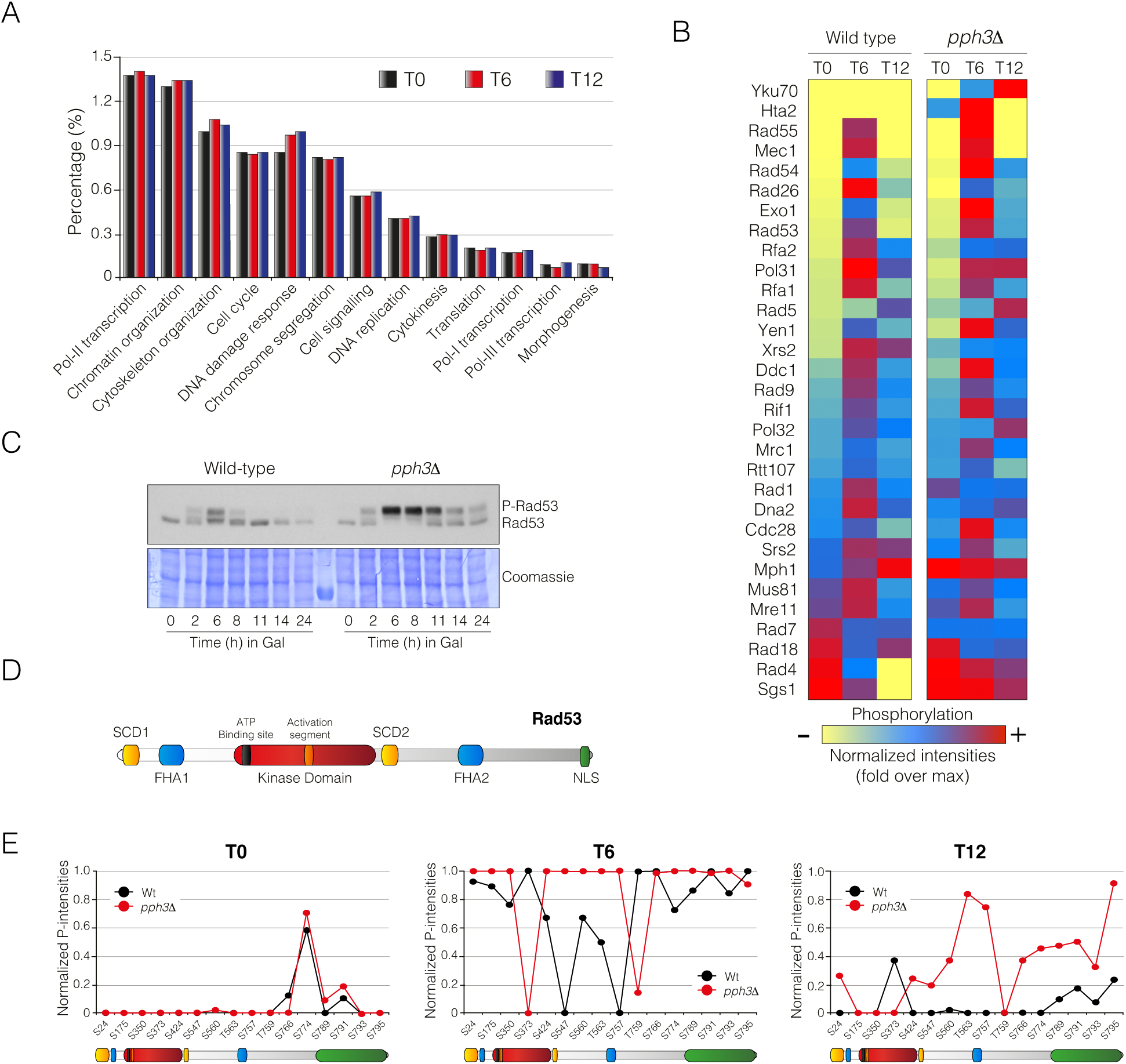
PP4 controls the phosphorylation state of multiple targets during the damage response. A) Identification of PP4 phospho-targets during the DNA damage response by mass spectrometry analysis. Wild-type and *pph3Δ* cells were grown overnight in raffinose-containing media prior the induction of the HO endonuclease by adding galactose to the culture. Samples from three independent experiments were taken at 0, 6 and 12 hours after the induction, processed and subjected to mass spectrometry. After isolation of PP4-dependent phospho-enriched peptides, their corresponded proteins were grouped into broad categories depending on their molecular function. Note that the main differences in peptide phospho-enrichment along the damage response is particularly observed in the “DNA damage response” category, denoting the great importance of PP4 activity during the response to DNA damage. B) Proteins containing at least one phospho-peptide enriched in the absence of PP4 within the category of “DNA damage response” were classified depending on their averaged phosphorylation. A classification containing the most relevant factors identified along the damage response is shown. Yellow, blue and red indicate relative amount protein phosphorylation (yellow, low; blue, medium; red, high). C) PP4 activity is required to maintain low levels of Rad53 phosphorylation during the response to DNA damage. Both wild-type and *pph3Δ* cells were grown overnight in raffinose-containing media and supplemented with galactose to induce HO expression. Samples were collected at the indicated time points. Proteins were TCA extracted and subjected to Western blotting. Coomassie staining is shown as loading control. D) Schematic representation of *S. cerevisiae* Rad53 illustrating the SQ/TQ clusters (yellow), forkhead-associated domains (blue), kinase domain (red) and nuclear localization signal (green). E) PP4 targets the FHA2 and NLS domains of Rad53. Graphs represent normalized P-intensities of the Rad53 phospho-peptides identified at 0, 6 and 12 hours from the HO induction.

Among the targets found, we focused our attention on the central checkpoint kinase Rad53. It has been proposed that Rad53 dephosphorylation by PP4 is needed for efficient cell cycle recovery from MMS treatment (O’Neill et al., 2007; Szyjka et al., 2008). However, the fact that Rad53 was already hyperphosphorylated during the repair stage prompted us to investigate whether PP4 could be also targeting Rad53 during the repair process. To validate the phosphatase activity of PP4 over Rad53 during the repair pathway, we used Western blotting to follow PP4-dependent changes in Rad53 phosphorylation after inducing a DSB in the YMV80 background used for the MS analysis. Confirming our previous observations, elimination of PP4 activity rendered Rad53 to an increased phosphorylation state when compared to the wild-type strain from 6 hours after the generation of the DSB (Fig. 2C). Rad53 consists of a central serine/threonine kinase domain, surrounded by two forkhead-associated (FHA) domains (FHA1 and FHA2) (Durocher et al., 1999) and two SQ/TQ cluster regions (SCD1 and SCD2) enriched in Mec1/Tel1-target phosphorylation sites (Traven and Heierhorst, 2005). Additionally, Rad53 includes a bipartite NLS domain at the C-terminal region (Smolka et al., 2005) required for its translocation into the nucleus (Fig. 2D). To gain insight into the specific PP4-dependent Rad53 dephosphorylation profile, we examined the MS data looking for changes in the phosphorylation pattern of the kinase in both wild-type and *pph3Δ* cells (Fig. 2E). We detected 16 residues in the Rad53 sequence that were potentially phosphorylated in response to DNA damage. A perfect match was found in the phosphorylation profile between a wild-type strain and a *pph3Δ* mutant at T0, with the majority of these sites unphosphorylated except for S774, a residue that has been shown to be a target of the Cdk in an unperturbed cell cycle (Diani et al., 2009). It has been reported that S774 phosphorylation does not play any role in the DDR but in morphogenesis (Diani et al., 2009). As expected, 6 hours after the expression of the HO endonuclease, a complete hyper-phosphorylation profile was clearly visible in both wild-type and *pph3Δ* strains. However, while a *pph3Δ* mutant presented a maximum level of phosphorylation around the FHA2 domain, wild-type cells displayed a more modest phosphorylation state at this region of the protein. After 12 hours from the DSB induction, all phosphorylation sites had been drastically reduced in the wild-type strain whereas *pph3Δ* cells still accumulated high levels of phosphorylation around the FHA2 domain and the NLS. Taken into account that phosphorylation of the FHA2 domain has been directly linked to a Rad53 active state (Sweeney et al., 2005) and that both FHA2 and NLS have been shown to be auto-Rad53 phosphorylation clusters (Chen et al., 2014; Smolka et al., 2005), these results clearly suggest that PP4 reduces Rad53 activity by counteracting its own auto-phosphorylation. Overall, these data demonstrate that PP4 is controlling the steady-state phosphorylation of multiple DNA damage factors during the repair of a DNA lesion, possibly by modulating the activity of Rad53 along the process. Importantly, the fact that PP4 activity over Rad53 takes place throughout the repair process indicates that this regulation might be central in the restoration of a DNA lesion.

### Restoration of Rad53 phosphorylation alleviates PP4-dependent defects in DNA repair

We have previously shown that in the absence of Pph3, the efficiency of DNA repair by SSA/BIR was affected when compared to a wild-type strain. In order to determine whether this reduction was directly linked to the high levels of Rad53 phosphorylation detected in the DDR, we analysed the effect of reducing Rad53 phospho-levels in *pph3Δ* cells during the execution of SSA/BIR pathways. It has been previously described that Rad9 binds the FHA2 domain of Rad53 to stimulate its phosphorylation by Mec1 in response to DNA damage (Sweeney et al., 2005). Accordingly, elimination of *RAD9* drastically reduced the high levels of Rad53 phosphorylation observed in the absence of Pph3 (Fig. 3A). In agreement with a negative effect of Rad53 hyper-phosphorylation over the repair efficiency, a double *pph3Δ rad9*Δ mutant was more efficient in repairing a DSB (Fig. 3B,D) and cells re-entered in the cell cycle faster than a single *pph3Δ* mutant (Fig. 3C). Accordingly, the lethality observed in *pph3Δ* cells growing on solid media containing 4-NQO, MMS, phleomycin and galactose was bypassed by ablating *RAD9* (Sup. Fig. 1A,C). Moreover, *PPH3* deletion in *rad9*Δ cells did not exacerbate the sensitivity of cells lacking *RAD9* to any of the genotoxic compounds tested (Sup. Fig. 1A,C), indicating that Pph3 acts in the same pathway of Rad9. These results confirm that the hyper-activation of Rad53 in the absence of PP4 activity affects DNA repair by SSA/BIR. It is important to note that while a double mutant *pph3Δ rad9*Δ retained some Rad53 phosphorylation (Fig. 3A), a single *rad9*Δ strain completely abolished its phosphorylation (Sup. Fig. 2A), suggesting that in the absence of Rad9, Rad53 retains certain capacity to be phosphorylated, but this phosphorylation is eliminated by the activity of PP4 during the DNA damage response. Moreover, a *rad9*Δ single mutant repaired more efficiently (Sup. Fig. 2B,D) and re-entered in the cell cycle with faster kinetics than the wild-type strain (Sup. Fig. 2C) confirming that Rad9 has a general effect in restraining SSA/BIR execution.

**Figure 3.**
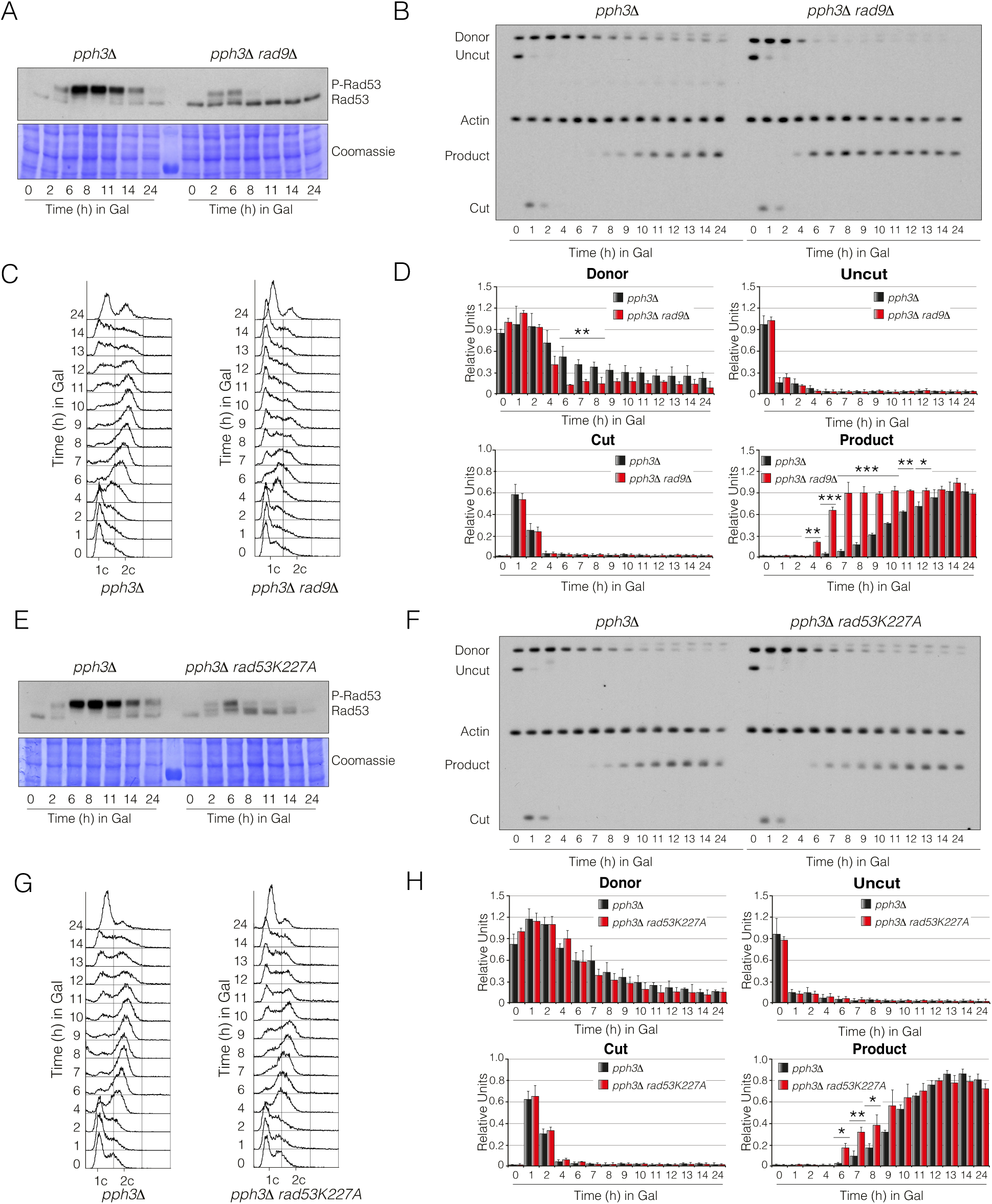
Reduction of Rad53 phosphorylation bypasses the defects in DNA repair observed in the absence of PP4 activity. A) Lack of Rad9 reduces the hyper-phosphorylation state of Rad53 observed in PP4-deficient cells. *pph3Δ* and *pph3Δ rad9*Δ cells were cultured overnight in media with raffinose and transfered to galactose-containing media to generate an HO-induced DSB. Samples were collected at the indicated time points, TCA extracted and subjected to Western blotting. Coomassie staining is shown as loading control. B) Elimination of Rad9 improves the repair efficiency of PP4-deficient cells. Physical analysis of *pph3Δ* and *pph3Δ rad9*Δ cells harbouring the DNA repair assay portrayed in figure 1A. Cells were grown in YP-Raffinose and transferred to media supplemented with galactose. Samples were collected at different time points, genomic DNA was extracted, digested with *Kpn*I and analysed by Southern blot. Blots were probed with an *U2* and *ACT1* (as loading control) DNA sequences. C) FACS profile for DNA content of samples taken from the experiment depicted in B). D) Graphs showing the quantification of the band signals obtained in the Southern blot illustrated in B). All data were normalized using the actin signal. Graphs represent the mean ± SD from three independent experiments. Replicates were averaged and statistical significance of differences assessed by a two-tailed unpaired Student’s t-test. E) The Rad53 hyper-phosphorylation state observed in the absence of PP4 derives from its own auto-phosphorylation activity. Cells from *pph3Δ* and *pph3Δ rad53K227A* strains were grown overnight in raffinose-containing media and galactose was added to induce a single HO DSB. Samples were collected at the indicated time points and subjected to Western blotting. Coomassie staining is shown as loading control. F) Elimination of Rad53 activity enhances DNA repair in the absence of PP4 activity. Southern blots of *pph3Δ* and *pph3Δ rad53K227A* cells carrying the DNA repair assay illustrated in figure 1A. Asynchronous cells growing in media supplemented with raffinose were transferred into galactose-containing media and samples were taken at the indicated time points. DNA was extracted, digested with *Kpn*I and blotted. Blots were probed with an *U2* DNA sequence and *ACT1* as a loading control. G) FACS profile for DNA content of samples collected from the experiment showed in F). H) Graphs representing the quantification of the band signals from the Southern blot experiment depicted in F). All data were normalized using the actin signal. Graphs represent the mean ± SD from three independent experiments. P-values were calculated using a two-tailed unpaired Student’s t-test.

Even though Rad53 phosphorylation directly depends on Rad9 activity, it has also been established that Rad9 is able to restrain resection by directly interfering with the activity of the Sgs1/Dna2 pathway (Bonetti et al., 2015; Ferrari et al., 2015; Gobbini et al., 2015), a function that could explain the improvement in DNA repair detected. Thus, we wondered if the exclusive down-regulation of the Rad53 hyper-phosphorylation levels observed in PP4-deficient cells might be sufficient to enhance SSA/BIR repair. We have previously shown that PP4 is specifically targeting the FHA2 domain of Rad53 in order to restrain its auto-phosphorylation during the DNA damage response (Fig. 2E). Supporting this observation, the substitution of the endogenous *RAD53* allele for a kinase dead *rad53K227A* drastically reduced the high levels of Rad53 phosphorylation observed in *pph3Δ* cells (Fig. 3E). This indicates that the Rad53 hyper-phosphorylation observed in the absence of PP4 comes from its own auto-phosphorylation activity. Besides, there were no apparent differences in the Rad53 phosphorylation status and DNA repair efficiency between a double *pph3Δ rad53K227A* and a single *rad53K227A* (Fig. 3E,F and Sup. Fig. 2E,F), confirming again that PP4 is specifically targeting Rad53 auto-phosphorylation sites to enhance DNA repair. Interestingly, Rad53 auto-activation is important for repairing by SSA/BIR, since the substitution of the wild-type version of RAD53 for the *rad53K227A* variant reduced the repair efficiency (Sup. Fig. 2F,G,H). Even with this counter-effect, the incorporation of the *rad53K227A* allele into the *pph3Δ* background slightly increased the capacity of the cells to repair a DNA lesion by SSA/BIR when compared to single *pph3Δ* mutants, and stimulated cell cycle re-entry once the DNA lesion had been restored (Fig. 3,F,G,H). Accordingly, the incorporation of the *rad53K227A* allele into a *pph3Δ* background increased cell viability on solid media containing MMS, phleomycin or galactose (Sup. Fig. 1B,C) supporting the idea of a toxic hyper-phosphorylation state of Rad53 in the absence of PP4 activity. Additionally, both *rad53K227A* and *pph3Δ rad53K227A* strains grew similarly on media containing most of the DNA-damaging agents tested, suggesting that both Rad53 and PP4 act in parallel in the same pathway. Thus, we conclude that PP4 participates in DNA repair by counterbalancing Rad53 activity during the DDR to allow the efficient execution of SSA/BIR repair pathways.

### PP4 is required for DNA end resection

We have previously shown that PP4 activity is required for maintaining a steady-state level of Rad53 phosphorylation during the DNA damage response, a function that is essential to promote an efficient repair by SSA/BIR. It has been proposed that phosphorylation of Exo1 by Rad53 is involved in a negative feedback loop that limits ssDNA accumulation during the resection process (Morin et al., 2008). On the other hand, Rad9 recruitment to the DSB vicinity depends on Rad53 activity (Gobbini et al., 2015). Importantly, Rad9 limits the resection activity of Sgs1/Dna2 by inhibiting the binding of Sgs1 to the DNA lesion (Bonetti et al., 2015; Ferrari et al., 2015). With all this information, we decided to check whether the hyper-phosphorylation levels of Rad53 detected in the absence of PP4 activity could affect SSA/BIR by restraining DNA end resection. To test for a role of PP4 in resection we used a JKM139 derivative strain containing a non-reparable HO system due to the elimination of both *HMR/HML* loci (Lee et al., 1998). As the 5’strand is degraded, restriction enzymes located in these areas are unable to cleave the generated ssDNA and, therefore, the intensity of the bands corresponding to the DNA fragments become diminished (Fig. 4A). First, we asked if the activation of a single non-reparable DSB in the absence of Pph3 also increased the levels of Rad53 phosphorylation during the DNA damage response. The fact that the induction of a non-reparable HO cut also led to an hyper-phosphorylated Rad53 state confirms that PP4 is acting over Rad53 from the initial steps of the repair process (Fig. 4B). To check for resection efficiency, we blocked the cells in G1 with alpha factor and released them into new media without the pheromone but containing galactose to induce the HO-break. As expected, while both strains entered the cell cycle and blocked in G2/M with the same kinetics by FACS, a *pph3Δ* mutant displayed a reduction in resection when compared to the wild-type (Fig. 4C). This indicates that lack of PP4 activity restrains SSA/BIR efficiency by interfering with resection at the DNA lesion.

**Figure 4.**
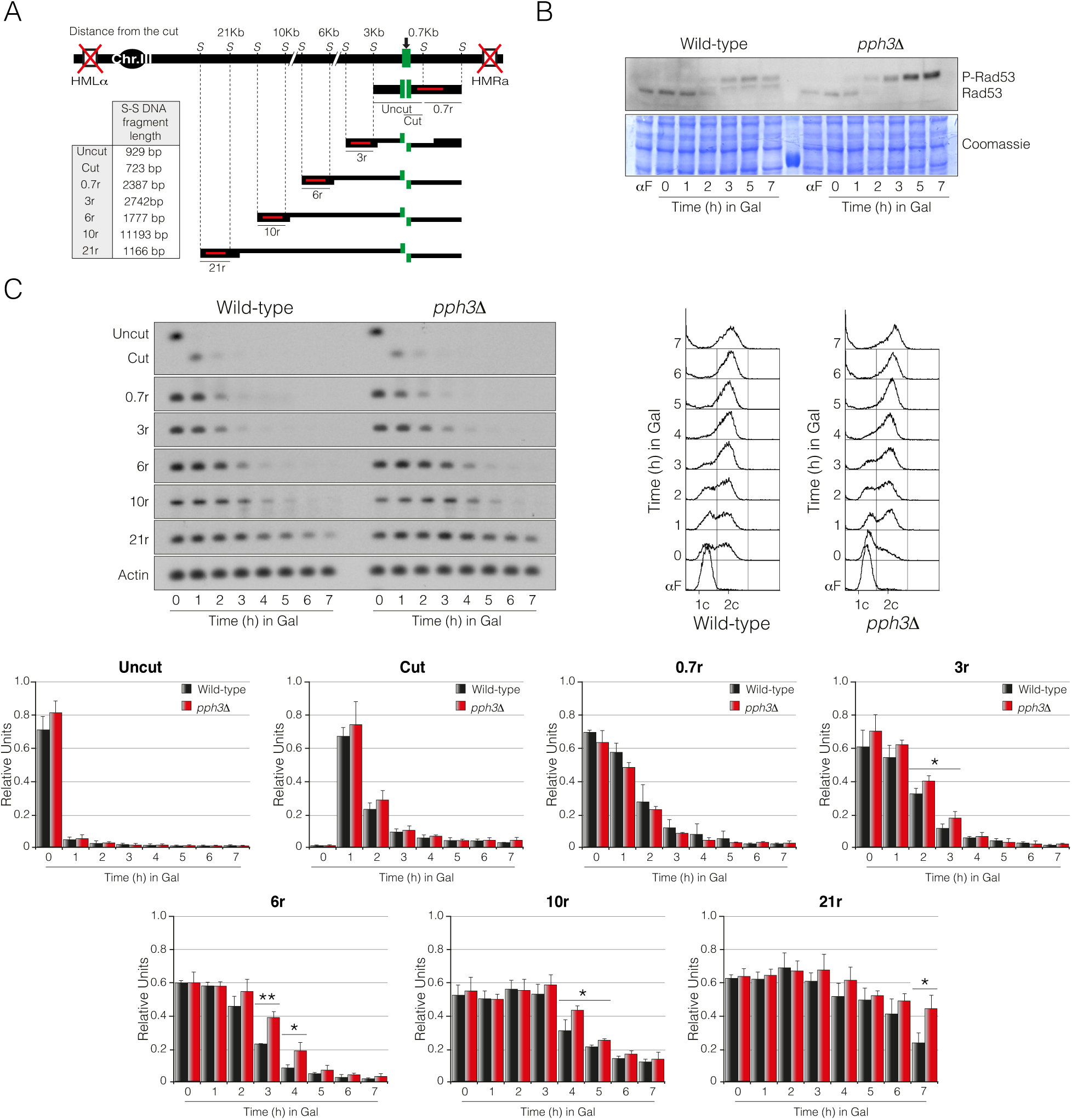
PP4 activity is required to stimulate DNA end resection. A) Schematic representation of the assay used to determine resection efficiency at the *MAT* locus. The diagram includes the recognition sites for *Sty*I and the localization of the probes used (red lines) for each distance. Table includes the *Sty*I*-Sty*I DNA fragments length for each probe. B) Rad53 is hyper-phosphorylated in the absence of PP4 activity during the induction of a non-reparable DSB. Exponentially wild-type and *pph3Δ* cells growing in YP-Raffinose were synchronized in G1 by using the α-factor pheromone and released into fresh media for 1 hour. Induction of HO expression was attained by adding galactose to the media and samples were taken at the indicated time points. Proteins were TCA extracted and subjected to Western blotting. Coomassie staining is shown as loading control. C) PP4 is required for an optional DNA end resection efficiency. Physical analysis by Southern blot of wild-type and *pph3Δ* strains containing the resection assay described in A). Samples were taken under the same conditions as in B), DNA extracted, digested with *Sty*I and blotted. A FACS analysis of the experiment is included. Band intensities were measured, normalized against actin and plotted. Graphs represent the mean ± SD from three independent experiments. P-values were calculated using a two-tailed unpaired Student’s t-test.

The helicase activity of Sgs1 is essential for the nuclease Dna2 to degrade the unwound ssDNA in a parallel pathway that involves the action of the exonuclease Exo1 (Zhu et al., 2008). Thus we wondered if PP4-dependent activation of resection relies on Dna2 and/or Exo1. To check for a possible function of PP4 in regulating the Exo1 pathway, we generated a single *exo1*Δ and a double *exo1*Δ *pph3Δ* mutant, and determined their influence in resection. The kinetics of DNA resection in an *exo1*Δ mutant was reduced when compared to a *pph3Δ* background. Moreover, lack of Exo1 in a *pph3Δ* mutant exacerbated the defect in resection (Fig. 5A). A similar effect was also observed when monitoring the resection intermediates generated during the induction of the unrepairable HO-cut (Sup. Fig. 3A). It is important to note that elimination of Exo1 did not affect the phosphorylation state of Rad53 (Sup. Fig. 3B), ruling out the possibility of Exo1 acting in the control of resection by interfering with Rad53 phosphorylation. These results demonstrate that the control of PP4 in DNA end resection is mainly carried out independently of Exo1. To investigate whether PP4 could be modulating the Dna2 resection pathway, we generated single *sgs1*Δ and double *sgs1Δ pph3Δ* mutants and determined their resection capacities. While a single *sgs1Δ* mutant resected with slower kinetics when compared to a *pph3Δ* mutant, a double *sgs1Δ pph3Δ* resected similarly than a single *sgs1Δ* (Fig. 5B). Only a minor defect was observed when looking at the resection intermediates between a single *sgs1*Δ and a double *sgs1Δ pph3Δ* mutant (Sup. Fig. 3C), a defect that might be attributed to a minor role of PP4 in controlling the Exo1 pathway. As previously reported, elimination of Sgs1 restrains the kinetics of Rad53 phosphorylation (Sup. Fig. 3D), probably due to its defects in DNA end resection and checkpoint activation (Gravel et al., 2008; Hegnauer et al., 2012). However, lack of Sgs1 did not influence the levels of Rad53 phosphorylation observed in *pph3Δ* cells (Sup. Fig. 3D) confirming that Sgs1 is not acting in the control of resection by interfering with Rad53 phosphorylation in the absence of PP4. These results demonstrate that PP4 controls DNA end resection mainly by regulating the activity of the Sgs1/Dna2 route, while a secondary and less robust mechanism comprise the modulation of the Exo1 pathway.

**Figure 5.**
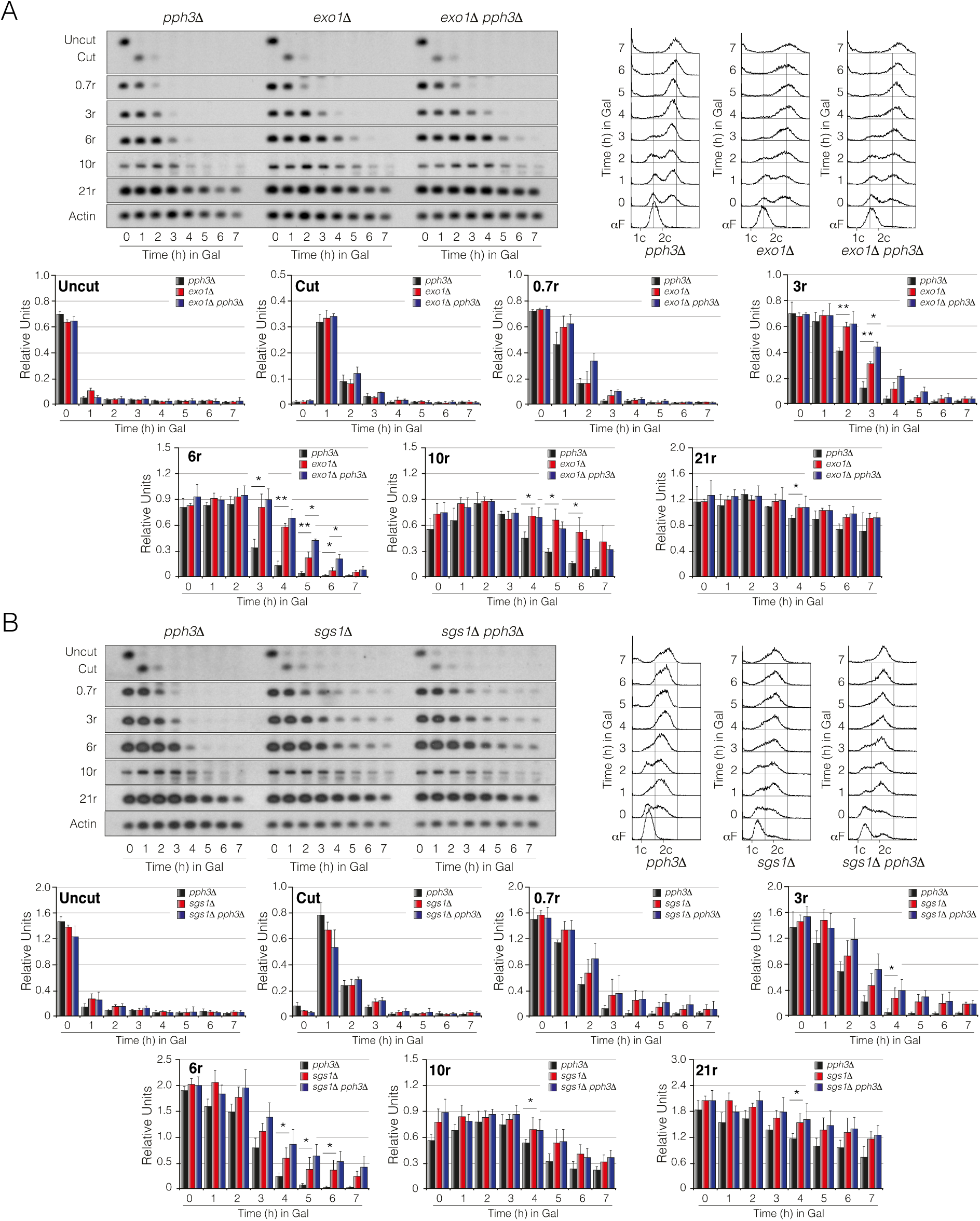
PP4’s role in DNA end resection is mainly driven by the modulation of the Sgs1/Dna2 pathway. A) Southern blots of *pph3Δ*, *exo1*Δ and *exo1*Δ *pph3*Δ cells grown in YP-Raffinose, blocked in G1 and released into fresh media containing galactose. Samples were taken at the indicated time points, DNA extracted, digested with *Sty*I and blotted. A FACS profile for DNA content is shown. The density of all bands was measured, normalized to actin and plotted. Graphs represent the mean ± SD from three independent experiments. P-values were calculated using a two-tailed unpaired Student’s t-test. B) Physical analysis by Southern blot of *pph3Δ*, *sgs1*Δ and *sgs1*Δ *pph3*Δ cells cultured in raffinose containing media, arrested in G1 and released into fresh media containing galactose. Samples were taken at the indicated time points, DNA extracted, digested with *Sty*I and blotted. A FACS profile for DNA content is included. The intensity of the bands was measured and charted. Graphs represent the mean ± SD from three independent experiments. P-values were calculated using a two-tailed unpaired Student’s t-test.

### Down-regulation of Rad53 phosphorylation suppresses the resection defects of cells lacking PP4

We have previously shown that *pph3Δ* mutants are impaired in executing DNA end resection in a pathway mainly dependent on Sgs1/Dna2. If this defect is due to the hyper-phosphorylation state of Rad53 observed in the absence of PP4 activity, then the reduction of its phosphorylation levels should bypass the resection deficiency of *pph3Δ* cells. Since Rad9 is required to promote Rad53 phosphorylation, we analysed the resection efficiency of a single *pph3Δ* mutant and in combination with a *RAD9* deletion. First, we confirmed that a double mutant *pph3Δ rad9Δ* reduced the high levels of Rad53 phosphorylation observed in a single *pph3Δ* (Fig. 6A). As expected, the kinetics of DNA resection in *pph3Δ rad9Δ* cells is markedly increased when compared to *pph3Δ* cells, indicating that *rad9Δ* suppresses the resection defect caused by the lack of Pph3 (Fig. 6A). It is important to note that due to the role of Rad9 in DNA damage checkpoint activation (Weinert and Hartwell, 1988), a double mutant *pph3Δ rad9Δ* is not able to completely block in G2/M, measured by FACS analysis after the induction of the HO break (Fig. 6A).

**Figure 6.**
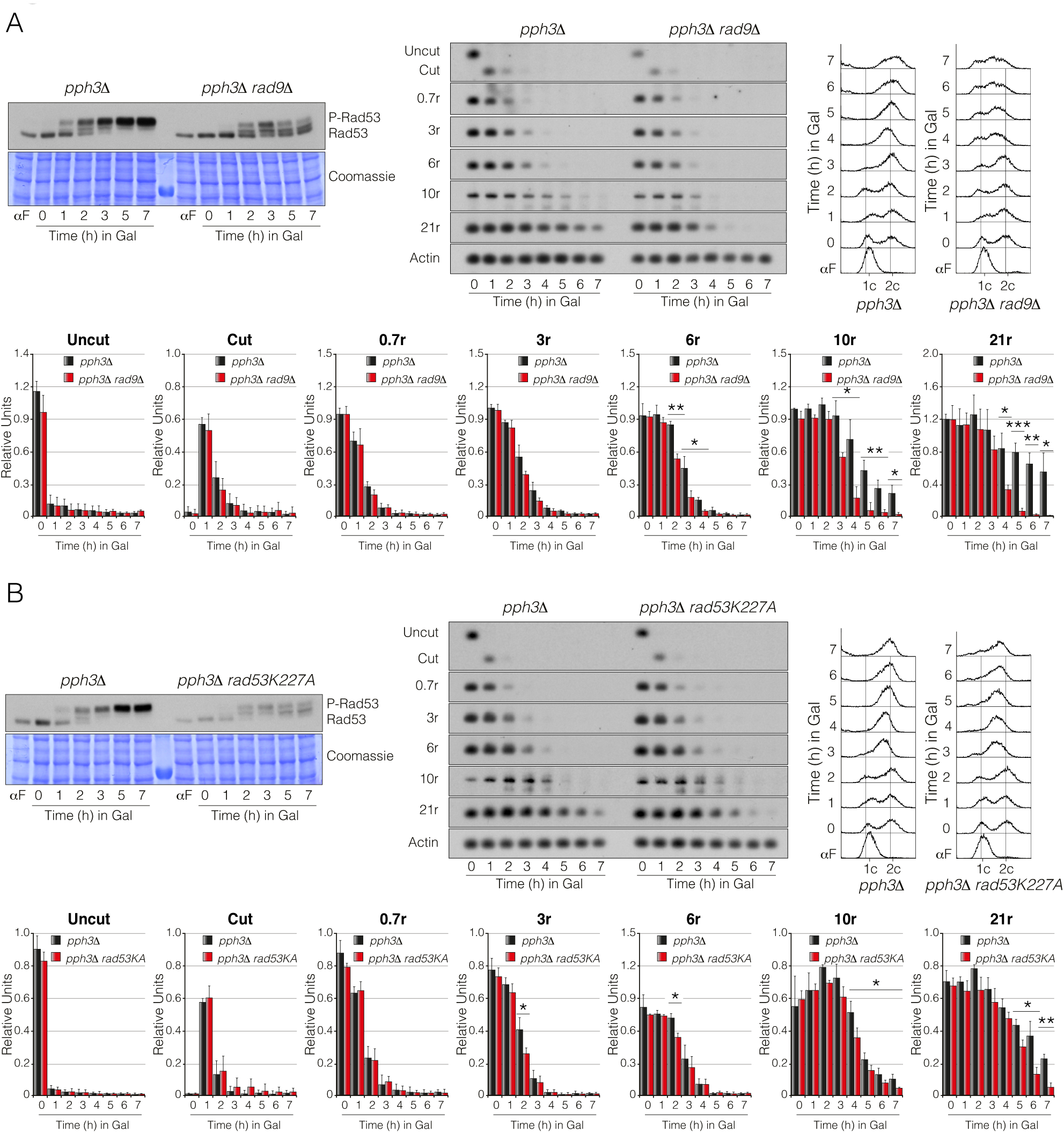
Down-regulation of Rad53 phosphorylation bypassed the resection defects observed in the absence of PP4 activity. A) Exponentially growing cells of *pph3Δ* and *pph3*Δ *rad9*Δ strains were synchronized in G1 by using the pheromone α-factor and released into fresh media for 1h. After inducing the HO expression by adding galactose to the media, samples were collected at the indicated time points for Western blotting, Southern blotting and FACS. Bands from the Southern blot were quantified, normalized against actin and plotted. Graphs represent the mean ± SD from three independent experiments. Replicates were averaged and statistical significance of differences assessed by a two-tailed unpaired Student’s t-test. B) Cells from *pph3Δ* and *pph3*Δ *rad53K227A* backgrounds were subjected to the same experimental conditions stated in A) and samples were processed for Western blotting, Southern blotting and FACS analysis. Band signals from the Southern blot were quantified, normalized against actin and charted. Graphs represent the mean ± SD from three independent experiments. P-values were calculated using a two-tailed unpaired Student’s t-test.

Even though elimination of Rad9 overcomes the high levels of Rad53 phosphorylation observed in a *pph3Δ* mutant, we cannot rule out that the improvement in DNA resection observed in a *pph3Δ rad9*Δ mutant could be also influenced by the direct role of Rad9 in counteracting Dna2 binding at the cut site (Bonetti et al., 2015; Ferrari et al., 2015). Thus we wondered if the exclusive reduction of Rad53 auto-phosphorylation might be sufficient to revert the resection defects observed in *pph3Δ* cells. As expected, introduction of a *rad53K227A* allele in a *pph3Δ* background drastically reduced the high levels of Rad53 phosphorylation observed in a single *pph3Δ* mutant (Fig. 6B). Importantly, the kinetics of resection in a double *pph3Δ rad53K227A* was increased when compared to a *pph3Δ* control strain (Fig. 6B), demonstrating that PP4 counteracts Rad53 auto-phosphorylation to reduce its inhibitory effect over resection. It is important to remark that the incorporation of the *rad53K227A* version does not influence the DNA damage checkpoint arrest at G2/M measured by FACS (Fig. 6B), implying that the effect of PP4 in controlling Rad53 phosphorylation during the initial steps of the repair pathway is exclusively related to its capacity to modulate the resection activity.

### PP4-dependent resection is essential for an efficient execution of SSA

As mentioned before, the DSB flanked by direct repeats present in the YMV80 background can be repaired by either SSA or BIR. While SSA repair is strictly dependent on resection for rendering the homologous sequences for annealing with each other, BIR involves the formation of a recombination-dependent replication fork to copy all distal sequences. Thus, we wondered whether the defects observed in DNA end resection in the absence of PP4 could affect all types of recombinational DNA repair or, by contrast, might be a process specifically required for repairing by SSA. To check for the involvement of PP4 in inter-chromosomal recombination, we used a genetic background containing a *MATa* sequence in chromosome V that can be recognized by the HO endonuclease and repaired with an uncleavable *MATa-inc* sequence located on chromosome III (Ira et al., 2003). In this strain, the donor *HM* loci have been eliminated and the *MATa-inc* mutation renders the repair product insensitive to successive HO cleavages (Sup. Fig. 4A). By using this approach, a unique reparable DSB by gene conversion is generated which can occur either with or without an associated crossover (Mazon and Symington, 2013). As the restriction fragments have an altered size when a crossover is formed, these events can be detected during the repair process. First, we confirmed that Rad53 was also hyper-phosphorylated in the absence of PP4 in an inter-chromosomal repair system (Sup. Fig. 4B). Gene conversion with no associated crossover was the predominant form of repair in both wild-type and *pph3Δ* cells. However we could not detect significant differences in the kinetics of DNA repair between both strains or in the dynamics of cell cycle re-entry measured by FACS (Sup. Fig. 4C). We neither detected differences in the proportion of gene conversion associated with crossing-over between the wild-type and the *pph3Δ* mutant among cells that repaired the DNA lesion (Sup. Fig. 4C). These data confirm that PP4 activity is not required to promote ectopic recombination with homologous sequences and suggest that the resection defects observed in the absence of Pph3 exclusively affect DNA repair by SSA.

To demonstrate this hypothesis, we took advantage of the role of Rad51 in promoting strand annealing. It has been demonstrated that Rad51 is not required for SSA execution but its deletion drastically affects BIR (Ivanov et al., 1996). Thus we disrupted *RAD51* in the YMV80 background strain and re-evaluated the effect of PP4 in DNA repair by SSA (Fig. 7A). Elimination of BIR by depleting Rad51 prolonged Rad53 phosphorylation when compared to a wild-type strain along the HO induction (Fig. 7B and Fig. 2C). As expected, a double mutant *rad51Δ pph3Δ* drastically increased Rad53 phopho-levels during the induction of the DSB compared with a single *rad51Δ* mutant (Fig. 7B). Correlating with the extended levels of phopho-Rad53 observed in a *rad51Δ* mutant, we monitored a delay in product accumulation and cell cycle re-entry when comparing to the wild-type strain (Fig. 7C and Fig. 1B), probably due to the lack of BIR contribution to the overall kinetics of DSB repair. In agreement with the defect observed in DNA repair efficiency, cell viability of *rad51Δ* single mutants was affected in drop assays on the presence of 4-NQO, HU, MMS and phleomycin (Sup. Fig. 1A,B). A similar sensitivity was also observed when the HO-break was induced by platting the cells on galactose-containing media (Sup. Fig. 1C). Remarkably, a double mutant *rad51Δ pph3Δ* was severely affected in DNA repair by SSA (Fig. 7C). This effect was corroborated by the synthetic lethality observed when the double mutant was grown on 4-NQO, MMS, phleomycin and galactose (Sup. Fig. 1A,B,C), indicating that PP4 acts in a different pathway of Rad51 during the response to DNA damage. However, no effect was observed when combining *rad51Δ* and *pph3Δ* in the sensitivity to HU, again suggesting that PP4 is not involved in the cellular response to replication stress. In all, these results demonstrate that PP4 is particularly necessary for the execution of DNA repair pathways that highly rely on resection to efficiently restore the DNA molecule.

**Figure 7.**
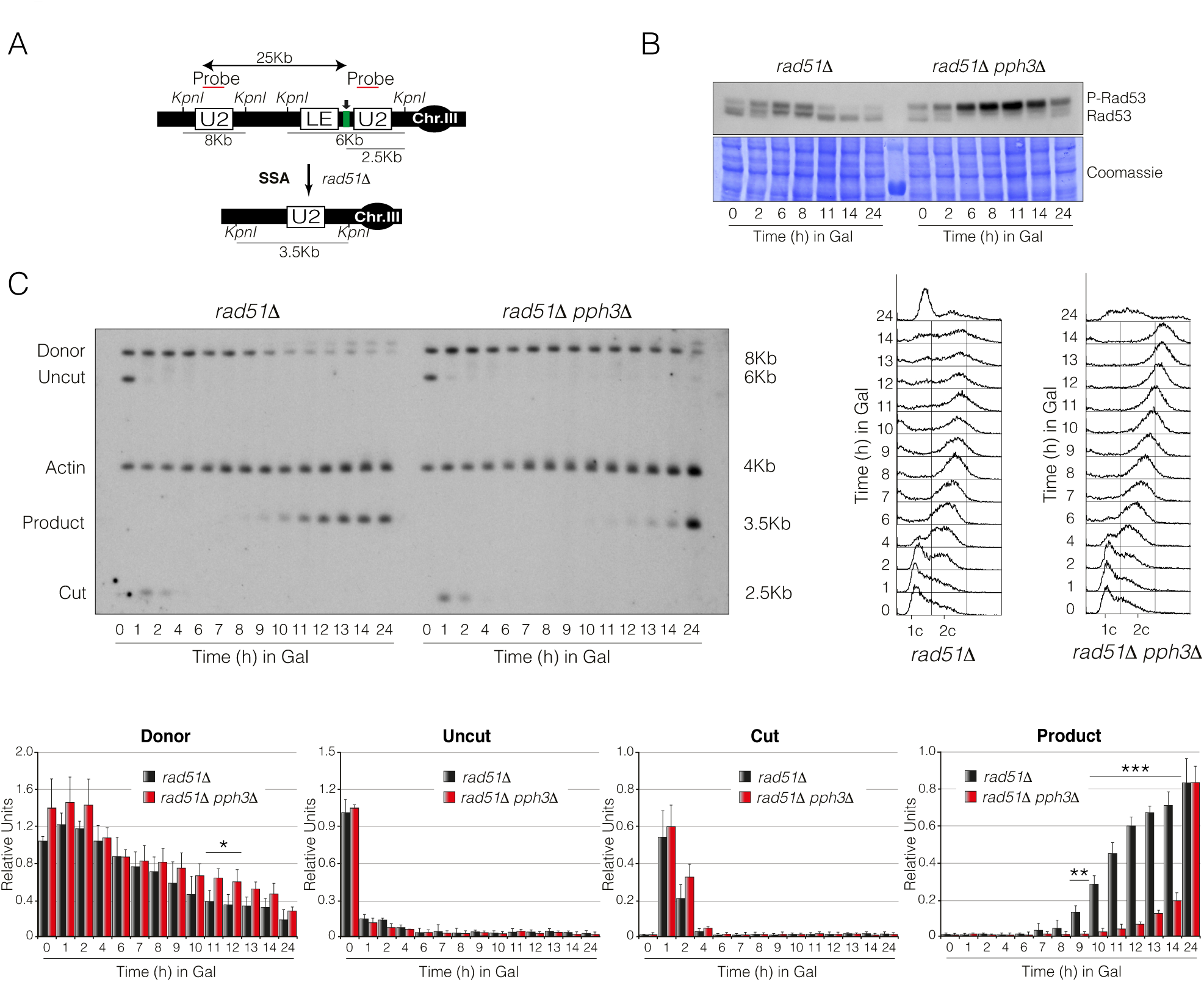
PP4’s function in DNA repair becomes essential when repairing by SSA. A) Schematic representation of the YMV80-based strain used to assess SSA efficiency. The localization of the *U2* probe, the restriction endonuclease cleavage sites used for Southern blot analysis and the expected bands size are indicated. Arrow depicts the localization of the HO break. B) PP4 activity is required for maintaining low levels of Rad53 phosphorylation during SSA execution. *rad51Δ* and *rad51Δ pph3Δ* cells were grown overnight in raffinose-containing media and supplemented with galactose to induce the expression of the HO. Samples were collected at the indicated time points. Proteins were TCA extracted and subjected to Western blotting. Coomassie staining is shown as loading control. C) Southern blot analysis of *rad51Δ* and *rad51Δ pph3Δ* strains carrying the DNA repair system depicted in A). Cells were grown overnight in YP-Raffinose before supplementing with galactose to induce HO expression, thus producing a DNA break at the *LEU2* locus on chromosome III. Samples were taken at time points indicated, genomic DNA was extracted, digested with *Kpn*I and analysed by Southern blot. Blots were hybridized with an *U2* probe DNA sequence and an *ACT1* probe was used as a loading control. FACS profiles for both strains are shown. Graphs represent the mean ± SD of the band signals from three independent experiments. All data were normalized using actin as a loading control. Replicates were averaged and statistical significance of differences assessed by a two-tailed unpaired Student’s t-test.

In order to confirm that the effect of PP4 over SSA was dependent on its ability to trigger DNA end resection, we determined the repair efficiency of the YMV80 background in a double *rad51Δ pph3Δ* and triple *rad51Δ pph3Δ rad9*Δ mutants. As expected, depletion of Rad9 in a *rad51Δ pph3Δ* strain drastically reduced Rad53 phosphorylation (Fig. 8A) nearly to wild-type levels (Fig. 2C). It is worth mentioning that while *rad51Δ pph3Δ rad9*Δ cells retained certain capacity to phosphorylate Rad53 (Fig. 8A), a *rad51Δ rad9*Δ strain completely abolished its phosphorylation (Sup. Fig. 5A), indicating that PP4 is actively dephosphorylating Rad53 during the repair of the DNA lesion by SSA. As expected, disruption of *RAD9* in a *rad51Δ pph3Δ* mutant completely restored SSA efficiency (Fig. 8B,D) and cells re-entered the cell cycle with a faster kinetic than the control strain (Fig. 8C). The improved repair efficiency was also corroborated by the increase in cell viability observed in the triple *rad51Δ pph3Δ rad9*Δ mutant compared to the double *rad51Δ pph3Δ* mutant on solid media containing 4-NQO, MMS, phleomycin and galactose (Sup. Fig. 1A,C). Likewise, elimination of *PPH3* in a *rad51Δ rad9*Δ strain did not exacerbate cell sensitivity when growing on the presence of all DNA-damaging agents used (Sup. Fig. 1A,C), demonstrating that Pph3 and Rad9 work in the same pathway. Importantly, deletion of *RAD9* in *rad51Δ* cells also improved SSA efficiency (Sup. Fig. 5B,D) and cell cycle re-entry (Sup. Fig. 5C), suggesting that the defects in SSA observed in the absence of Rad51 are directly linked to the negative effect that Rad9 exerts over resection.

**Figure 8.**
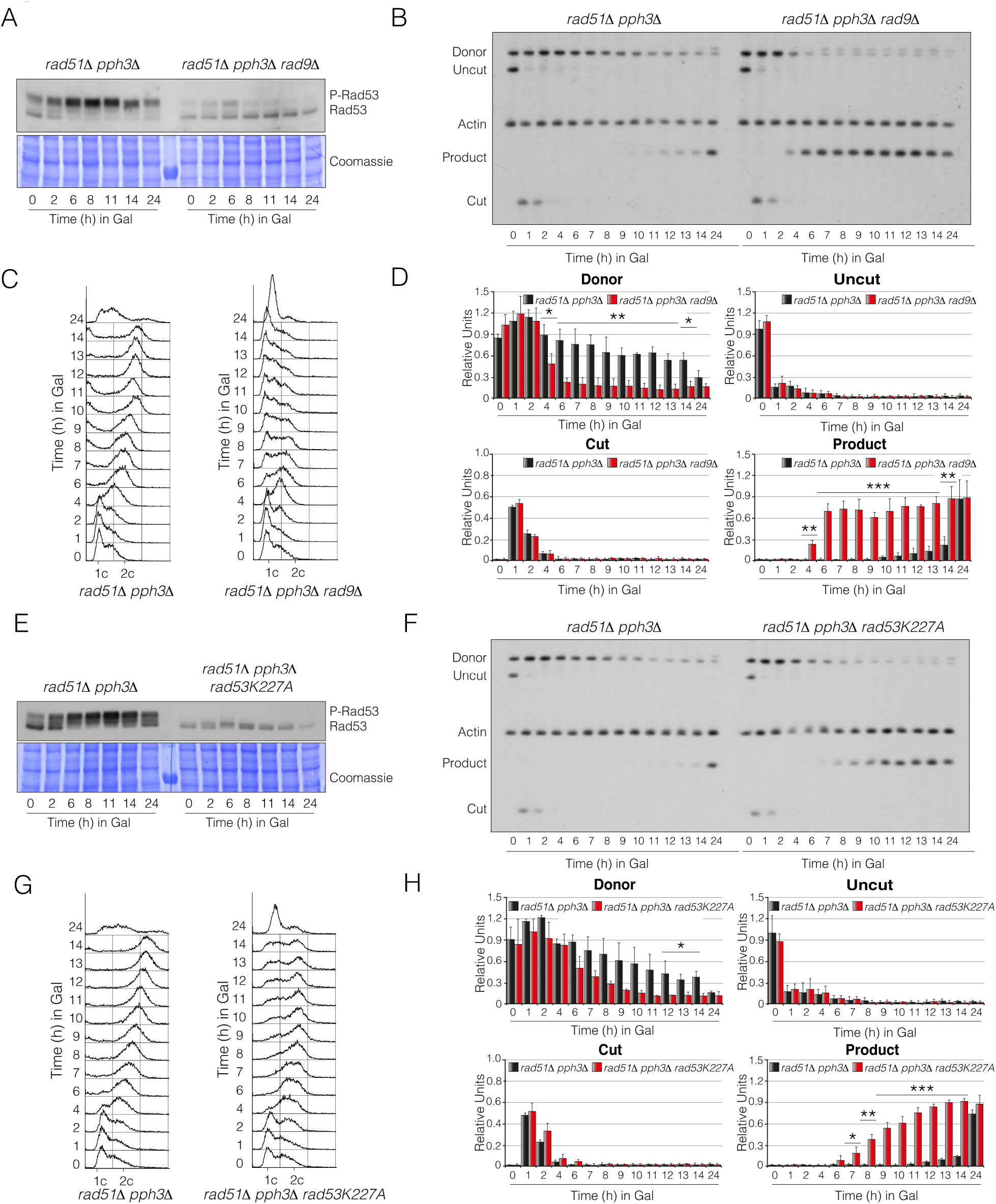
Reduction of Rad53 phospho-levels in PP4 –deficient cells improves DNA repair by SSA. A) Disruption of Rad9 reduces the hyper-phosphorylation state of Rad53 observed in cells lacking PP4 activity. Exponentially growing YP-Raffinose cell cultures of YMV80 derivative strains containing *rad51Δ pph3*Δ and *rad51Δ pph3Δ rad9*Δ deletions were supplemented with galactose to induce the expression of the HO endonuclease. Samples were collected at the indicated time points, TCA extracted and subjected to Western blotting. Coomassie staining is shown as loading control. B) Lack of Rad9 improves the repair efficiency of PP4-deficient cells by SSA. Physical analysis of *rad51Δ pph3*Δ and *rad51Δ pph3Δ rad9*Δ backgrounds harbouring the DNA repair assay portrayed in figure 7A. Cells were grown overnight in YP-Raffinose and supplemented with galactose. Samples were taken at different time points, genomic DNA extracted, digested with *Kpn*I and analysed by Southern blot. Blots were probed with an *U2* probe and *ACT1* was used as loading control. C) FACS profile for DNA content of samples collected from the experiment depicted in B). D) Graphs represent the quantification of the band signals obtained from the Southern blot depicted in B). All experiments were normalized using the actin signal. Graphs represent the mean ± SD from three independent experiments. Replicates were averaged and statistical significance of differences assessed by a two-tailed unpaired Student’s t-test. E) Rad53 hyper-phosphorylation state displayed in PP4-deficient cells is restored when introducing a kinase dead allele of *RAD53*. Cells from *rad51Δ pph3*Δ and *rad51Δ pph3Δ rad53K227A* background strains were grown overnight in media containing raffinose and galactose was added to induce a single HO cut. Samples were collected at the indicated time points and subjected to Western blotting. Coomassie staining is shown as loading control. F) Elimination of Rad53 activity enhances DNA repair by SSA in the absence of PP4 activity. Southern blot of *rad51Δ pph3*Δ and *rad51Δ pph3Δ rad53K227A* cells carrying the DNA repair system depicted in figure 7A. Exponentially growing YP-Raffinose cells were supplemented with galactose and samples were taken at the indicated time points. DNA was extracted, digested with *Kpn*I and blotted. Blots were probed with an *U2* DNA sequence and *ACT1* as loading control. G) FACS profile for DNA content of samples collected from experiment shown in F). H) Graphs representing the quantification of the band signals detected in the Southern blot illustrated in F). All data were normalized using the actin signal. Graphs represent the mean ± SD from three independent experiments. P-values were calculated using a two-tailed unpaired Student’s t-test.

To directly connect the ability of PP4 to counteract Rad53 auto-phosphorylation with SSA repair, we constructed a *rad51Δ pph3Δ* strain containing the catalytic-dead allele *rad53K227A*. Again, the incorporation of the kinase-dead version of Rad53 restored the high levels of Rad53 phosphorylation observed in *rad51Δ pph3Δ* cells (Fig. 8E), confirming that PP4 activity counterbalances Rad53 function during DNA repair. Remarkably, the integration of the *rad53K227A* allele in *rad51Δ pph3Δ* cells considerably improved their ability to repair by SSA (Fig. 8F,H) and thus, their proficiency to re-enter in the cell cycle (Fig. 8G). This was reinforced by the fact that the incorporation of the *rad53K227A* allele in a double *rad51Δ pph3Δ* background strain efficiently recovered cell viability in the presence of most of the DNA-damaging agents tested (Sup. Fig. 1B,C). However, only a mild enrichment in SSA efficiency was detected when introducing the *rad53K227A* allele in a *rad51Δ* control strain (Sup. Fig. 5E,F,G,H). Overall, these data demonstrate that PP4 targets Rad53 to restrain its own auto-phosphorylation in order to relieve the resection inhibition exerted by Rad9/Exo1 and elicit a proficient execution of the SSA repair pathway.

### PP4 controls the phosphorylation state of Rad9 and Exo1 by acting over Rad53

We have previously shown that PP4 enhances DNA repair by stimulating DNA end resection through its ability to restrain Rad53 auto-phosphorylation along the DNA damage response. This observation envisages that Rad53 hyper-activation in the absence of PP4 activity might influence the steady-state phosphorylation of downstream targets operating in the resection process. Supporting this hypothesis both Rad9 and Exo1 presented a differential distribution of several phospho-residues between a wild-type and a *pph3Δ* mutant in our MS experiment. Rad9 contains a Chk1 activation domain at the N-terminus (Abreu et al., 2013) close to a Mec1 serine cluster domain. The C-terminus includes a TUDOR domain involved in the binding of Rad9 to the DNA throughout its interaction with H3K79me (Grenon et al., 2007) and a protein-protein recognition module BRCT domain (Bork et al., 1997) (Fig. 9A). We detected 40 residues in Rad9 with the potential to be phosphorylated along the induction of a reparable HO break. A random distribution of phosphorylation was observed along the protein before the induction of the DNA lesion with a similar pattern between the wild-type and the *pph3Δ* mutant (Fig. 9B). After six hours from the induction of the HO endonuclease, a global increase in phosphorylation was observed in both wild-type and *pph3Δ* mutant. However, the absence of PP4 activity specifically increased the phosphorylation levels of several phospho-residues comprised between the serine cluster domain and the TUDOR domain (Fig. 9B). Considering that this domain is responsible for the binding of Rad9 to the DNA (Grenon et al., 2007) together with the fact that its association depends on Rad53 activity (Gobbini et al., 2015), it seems that the hyper-phosphorylated state of Rad53 in the absence of PP4 activity might be controlling DNA resection by modulating Rad9 affinity for the DNA lesion. After 12 hours from the HO induction, Rad9 phosphorylation levels decayed in both strains (Fig. 9B), indicating that other phosphatases are collaborating with PP4 in order to reduce Rad9 phosphorylation once the repair process has taken place. In order to validate these results we used Phos-Tag gels to follow PP4-dependent changes in the phosphorylation state of Rad9 after inducing a non-reparable DSB. As previously shown, upon induction of the DNA lesion Rad53 presented a higher phosphorylation state in the absence of PP4 activity than the parental strain, a situation that was reverted when the *rad53K227A* variant replaced the endogenous Rad53 (Fig. 9C). Ratifying the data obtained by the MS assay, Rad9 was already phosphorylated before the induction of the HO. However, after inducing the DNA break, while we observed accumulation of higher molecular weight bands that denoted a general hyper-phosphorylation state, a few low molecular weight forms were also detected (Fig. 9D). These fast migrating bands were only detected in the wild-type strain, but not in a *pph3Δ* strain (Fig. 9D), demonstrating that the lack of PP4 activity also affects the phosphorylation status of Rad9. Notoriously, the introduction of the *rad53K227A* allele drastically diminished the phosphorylation state of Rad9, demonstrating that Rad53 is in part responsible for the phosphorylation pattern observed in Rad9 during the DNA damage response (Fig. 9D).

**Figure 9.**
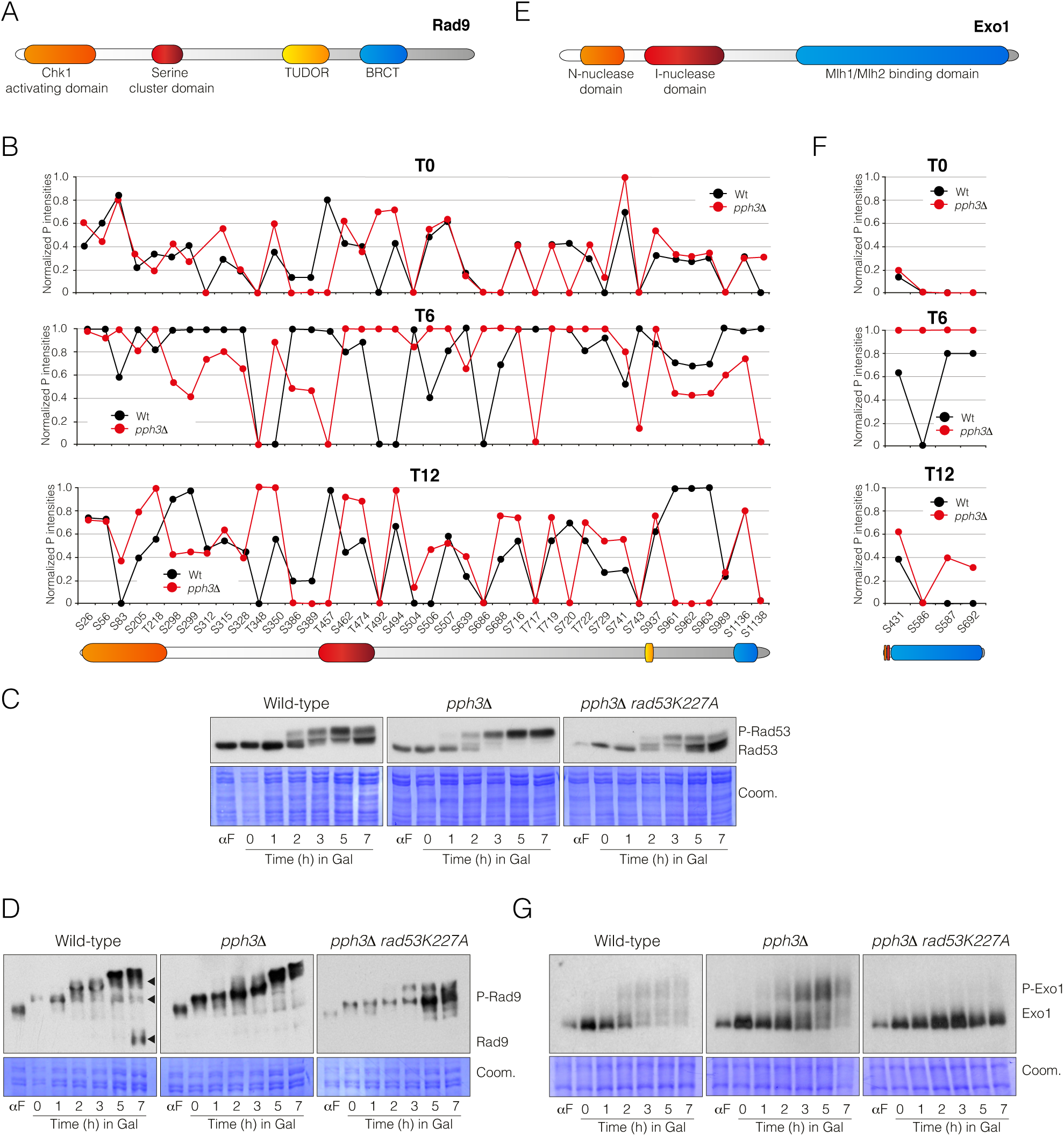
PP4 modulates Rad9 and Exo1 phosphorylation by acting over Rad53. A) Schematic representation of *S. cerevisiae* Rad9 illustrating the Chk1 activating domain (orange), a Mec1 serine cluster domain (red), a DNA interaction TUDOR domain (yellow) and a protein-protein recognition module BRCT domain (blue). B) Graphs representing normalized P-intensities of the Rad9 phospho-peptides identified by mass spectrometry at 0, 6 and 12 hours from the HO induction. C) Western blots of wild-type, *pph3Δ* and *pph3*Δ *rad53K227A* cells subjected to a non-reparable DSB. Cells were synchronized in G1 by using α-factor and released into fresh media containing galactose. Samples were taken at the indicated time points, TCA extracted and blotted using antibodies directed against Rad53. Coomassie staining is included as loading control. D) The steady-state phosphorylation of Rad9 is modulated by the effect of PP4 over Rad53. Wild-type, *pph3Δ* and *pph3*Δ *rad53K227A* cells containing the endogenous Rad9 tagged with the HA epitope were subjected to the same experimental conditions as in C) and blotted using HA antibodies. Coomassie staining is depicted as loading control. E) Diagram depicting *S. cerevisiae* Exo1, including its N and I nuclease domains (orange and red respectively) and its Mlh1/Mlh2 binding domain (blue). F) Schematic representation of normalized P-intensities of the Exo1 phospho-peptides found by mass spectrometry at 0, 6 and 12 hours from the induction of the HO endonuclease. G) Exo1 phosphorylation is regulated by the effect that PP4 exert over Rad53. Wild-type, *pph3Δ* and *pph3*Δ *rad53K227A* cells containing the endogenous Exo1 tagged with the MYC epitope were subjected to the same experimental conditions as in C) and blotted using MYC antibodies. Coomassie staining is depicted as loading control.

It has been postulated that Exo1 phosphorylation by Rad53 comprises a mechanism that limits the formation of resected DNA tracks during the damage response (Morin et al., 2008). Supporting this information Exo1, appeared hyper-phosphorylated in the absence of PP4 activity in our MS screening. The nuclease domain of Exo1 is located at the N-terminal one-third of the protein, while at the C-terminus resides a Mlh1/Mlh2 interacting domain (Dherin et al., 2009; Segurado and Diffley, 2008) (Fig. 9E). We could not detect any phosphorylated residue before induction of the HO break neither in the wild-type strain nor in the *pph3Δ* mutant (Fig. 9F). However, six hours after the generation of the DSB, we detected phosphorylation of four serine residues, all of them comprised at the Mlh1/Mlh2 binding domain. Interestingly, the absence of PP4 activity increased the average level of all phosphorylated residues compared to the wild-type (Fig. 9F). Notably, phosphorylation of Ser587 and Ser692 has been previously described to have a role in restraining Exo1 activity in response to uncapped telomeres and camptothecin treatment (Morin et al., 2008). After 12 hours from the HO induction, while a wild-type strain reduced the level of phosphorylation of all the residues detected, a *pph3Δ* mutant still contained high levels of phosphorylation at most of these sites (Fig. 9F). Confirming these data, Exo1 phosphorylation levels were increased in Phos-Tag gels in the absence of Pph3 (Fig. 9G). Hyper-phosphorylation of Exo1 was reverted when the endogenous Rad53 was substituted for the *rad53K227A* allele, confirming that PP4 controls Exo1 levels by reducing the levels of Rad53 phosphorylation in the presence of a DSB (Fig. 9G). Overall, these results demonstrate that in the absence of PP4 activity, the high level of Rad53 phosphorylation is responsible for the increase in Rad9 and Exo1 phosphorylation during the DNA damage response.

### The defects in DNA resection and repair observed in PP4-deficient cells are bypassed by eliminating Ser129 phosphorylation of histone H2A

We have previously shown that PP4 largely enhances DNA end resection by overcoming the inhibitory effect that Rad9 exerts over the Sgs1/Dna2 pathway (Fig. 5B and Fig. 6A). It has been postulated that Rad9-dependent inhibition of Sgs1/Dna2 is attained by its physical interaction at the DSB vicinity, a process that depends on Rad53 (Gobbini et al., 2015) and phosphorylation of histone H2A at Ser129 (Hammet et al., 2007; Toh et al., 2006). Surprisingly, PP4 inactivation increased Rad9 phosphorylation specifically at the TUDOR domain (Fig. 9B), suggesting that PP4-dependent Rad53 regulation might be vital to avoid excessive binding of Rad9 to the DNA break. To attain this question, we decided to determine whether elimination of S129 phosphorylation of H2A bypasses the phenotypes observed in the absence of PP4 activity. We constructed a strain in which S129 of H2A was substituted by a stop codon in both *HTA1* and *HTA2* alleles (*hta1/hta2-S129\**) (Downs et al., 2000) and determined its influence in rescuing both DNA end resection and repair by SSA/BIR in cells lacking *PPH3* by using a YMV80 strain. Elimination of S129 from H2A in a *pph3Δ* background did not affect Rad53 phosphorylation when compared with a *pph3Δ* mutant in response to a DSB (Fig. 10A). However, these cells considerably increased the repair efficiency by SSA/BIR (Fig. 10B,C) and re-entered in the cell cycle with faster kinetics (Fig. 10D). Since Rad53 was similarly phosphorylated in both strains, this result suggests that the high levels of Rad53 phosphorylation observed in the absence of PP4 are specifically modulating Rad9 capacity to interact with H2A phosphorylated at S129. Strikingly, a double mutant *pph3Δ hta1/hta2-S129\** resected faster than a single *pph3Δ* indicating that the improvement in DNA repair is linked to its ability to stimulate DNA resection (Fig. 10B,C) due to elimination of the negative effect that Rad9 exerts over the Sgs1/Dna2 pathway.

**Figure 10.**
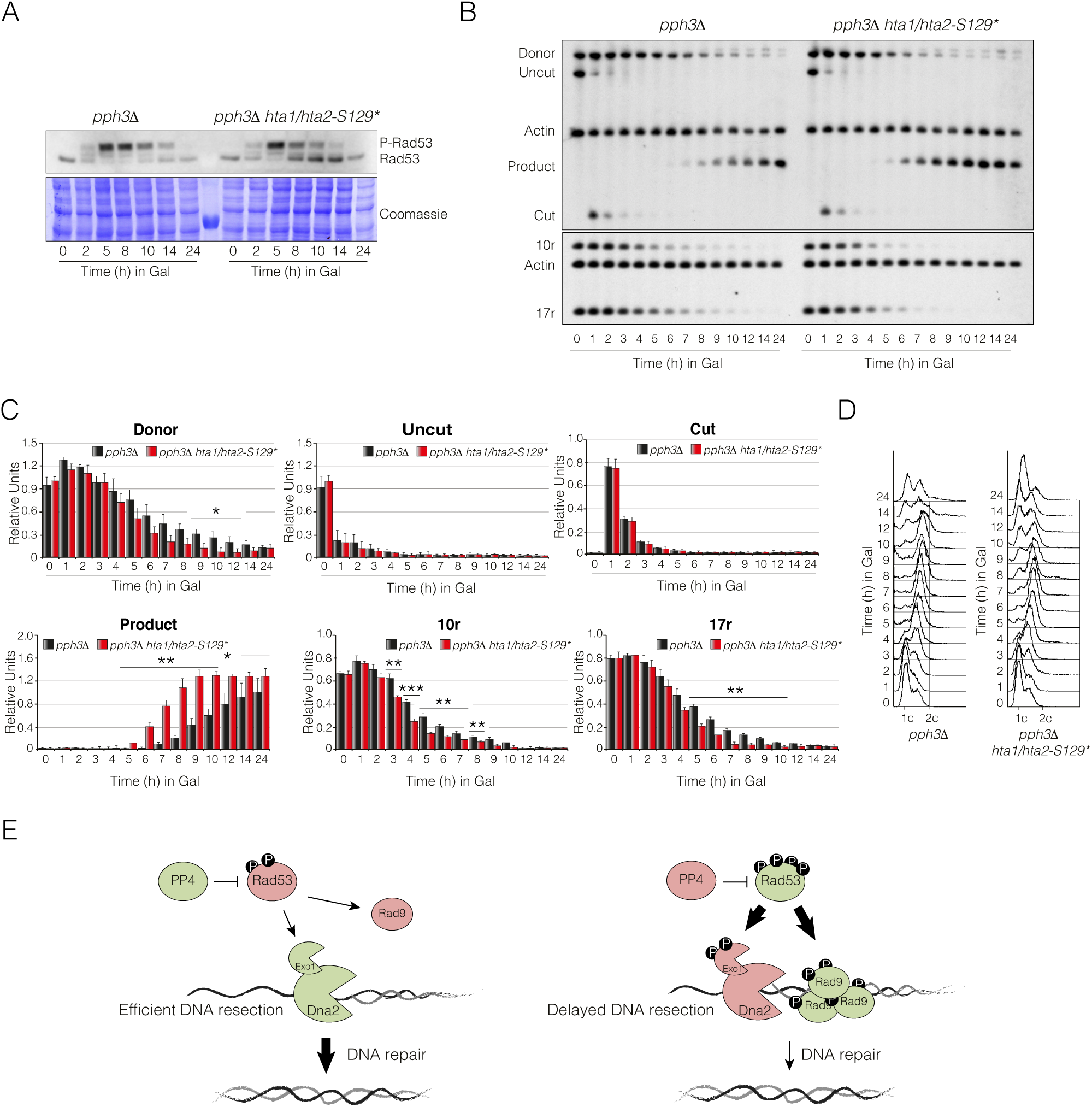
Elimination of H2A phosphorylation at Ser129 rescues both DNA end resection and repair defects of cells lacking PP4 activity. A) Removal of Ser129 phosphorylation from histone H2A does not influence in the Rad53 phosphorylation profile during the induction of a single DSB. YP-Raffinose cell cultures of YMV80 derivative *pph3Δ* and *pph3Δ hta1/hta2-S129\** cells were supplemented with galactose to induce the expression of the HO endonuclease and samples were collected at the indicated time points. Proteins were extracted by a TCA method and subjected to Western blotting. Coomassie staining is shown as loading control. B) Elimination of H2A phosphorylation at Ser129 improves the repair efficiency of PP4-deficient cells by SSA/BIR. Physical analysis of *pph3Δ* and *pph3Δ hta1/hta2-S129\** cells harbouring the DNA repair assay portrayed in figure 1A. Cells were grown overnight in YP-Raffinose and supplemented with galactose. Samples were taken at different time points, genomic DNA extracted, digested with *Kpn*I and analysed by Southern blot. Blots were probed with an *U2* probe and *ACT1* was used as loading control. For the analysis of resection efficiency, blots were hybridized with probes located at 10 Kb and 17 Kb upstream the HO cut site. C) Graphs represent the quantification of the band signals obtained from the Southern blot experiment shown in B). All data were normalized using the actin signal. Graphs show the mean ± SD from three independent experiments. Replicated were averaged and statistical significance of differences assessed by a two-tailed unpaired Student’s t-test. D) FACS profile for DNA content of samples collected in B). E) Model for the role of PP4 in resection activation. The activity of PP4 is required for maintaining low levels of Rad53 phosphorylation during the initial steps of the repair pathway (left panel). This steady-state of Rad53 phosphorylation attenuates its activity, a feature that affects the phosphorylation levels of down-stream targets as Exo1 and Rad9. An Exo1 dephosphorylation state increases its processivity, while Rad9 hypo-phosphorylation limits its presence at the DNA break, allowing a full activation of the Sgs1/Dna2 pathway. In the absence of PP4 activity (right panel), hyper-activated Rad53 triggers Exo1 and Rad9 hyper-phosphorylation. While Exo1 phosphorylation slightly affects resection, hyperphosphorylated Rad9 binds to chromatin and restrains DNA resection by disturbing Sgs1/Dna2 activity. Green and red denote active and inactive respectively.

Altogether, our results support a model whereby Rad53 dephosphorylation by PP4 stimulates DNA end resection during the initial steps of the repair pathway. The maintenance of a reduced Rad53 activity during this stage of the repair process ensures a low steady-state phosphorylation of multiple targets during the response, including Rad9 and Exo1 (Fig. 10E). Reduction in Rad9 phosphorylation counteracts its inhibitory function over the Sgs1/Dna2 complex by reducing its affinity to chromatin, ensuring a robust and efficient resection activation needed for repair mechanisms that rely on long ssDNA trails to ensure the success of the repair (Fig. 10E). Interestingly, these results reveal that a DNA damage checkpoint hyper-activation is deleterious for DNA repair and demonstrate that the maintenance of the steady-state phosphorylation of multiple components of the DDR ensures the execution of a proficient checkpoint activation while preserving the activity of the repair machinery. Remarkably, this novel concept envisages that kinases and phosphatases must cooperate in the repair of a DNA lesion in order to orchestrate the multiple and complex mechanisms enclosed in the DNA damage response.

## Discussion

Phosphorylation in response to DNA damage has been one of the main subjects in the field of DNA repair, focusing mainly in the regulation of the kinases operating in the DDR. Even though there is a consensus agreement that dephosphorylation is essential for restoring the effects imposed by the DDR-kinases, less is known about the role of protein phosphatases during the execution of the DNA damage response. To date, most of the functions attributed to the DDR-phosphatases during the response to a DNA lesion have been related to their capacity to inactivate the DNA damage checkpoint and promote cell cycle re-entry. However, taking into account the great complexity of the different pathways encompassed in the DDR and the multiple events taking place during the repair of a DNA abduct, it is tempting to speculate that the fine-tune phosphorylation along the different steps of the repair process might be essential to ensure the restoration of the DNA molecule. Supporting this theory, several DNA repair factors have multiple phospho-sites that are phosphorylated and dephosphorylated concurrently, and the kinetics are potentially regulated by kinases and phosphatases working in tandem. Accordingly, new studies have pointed forward the importance of several phosphatases in the DDR, not only by promoting cell cycle re-entry after the lesion has been fixed, but also by playing a direct role at the repair level.

In this study we have evaluated the importance of the PP4 phosphatase in the repair of a DNA lesion in the budding yeast *S. cerevisiae*. Elimination of PP4 activity rendered cells to a defect in DNA end resection that led to an impairment in the execution of homologous recombination pathways that require long-range resection for their accomplishment. Surprisingly, MS data of cells lacking PP4 activity revealed an elevated number of DNA damage factors with atypical high levels of phosphorylation along the damage response. This global DNA damage-dependent up-regulation of DDR components suggests a role of PP4 in controlling the activity of the main kinases operating in the damage response. Confirming this hypothesis, Rad53 was hyper-phosphorylated in response to a DSB in the absence of PP4 activity. This result, together with the observation that PP4 exhibited a preference for dephosphorylating Rad53 auto-phosphorylation sites, implies that the modulation of this kinase could influence the steady-state phosphorylation of multiple targets during the damage response. Still, we cannot discard that PP4 might also collaborate in the dephosphorylation of Rad53 targets during the response to a DNA lesion. However, the fact that some of the targets found to be hyper-phosphorylated in the absence of PP4 activity restored their phosphorylation levels when inhibiting Rad53 auto-phosphorylation claims for a main role of the phosphatase in directly controlling the phosphorylation status of the kinase. It is also important to note that PP4 might be controlling the activity of Rad53 by modulating the phosphorylation/activity of upstream kinases. In this regard, it has been recently shown that the absence of Mec1-S1991 phosphorylation in *mec1-100* cells is restored upon *PPH3* deletion, indicating that PP4 indeed dephosphorylates Mec1 (Hustedt et al., 2015). However, phosphorylation of this residue requires also the activity of Rad53 (Hustedt et al., 2015), suggesting that PP4 might be indirectly affecting Mec1 phosphorylation throughout the modulation of Rad53 activity. Nevertheless, we could not detect differences in Mec1 phosphorylation between wild-type and PP4-deficienct cells in our MS data during the induction of an HO-break, indicating that PP4 has not an important role in controlling the phosphorylation of Mec1 under these conditions.

How does PP4-dependent Rad53 dephosphorylation affect the execution of the DNA damage response? One puzzling observation extracted from our experiments is that, while PP4 is essential during DNA damage recovery when cells are treated with MMS (O’Neill et al., 2007), we did not observe a clear defect in cell cycle re-entry upon DNA repair. There are two non-mutually exclusive interpretations to explain this apparent controversy. One possibility is that PP4-dependent checkpoint deactivation depends on the extent of the damage infringed and/or the type of genotoxic stress. Accordingly, Pph3 is dispensable for Rad53 recovery after replication stress (Travesa et al., 2008). Moreover, the activity of Rad53 during MMS recovery is quite similar between wild-type and *pph3Δ* cells, indicating that Rad53 phosphorylation and kinase activity can be separated (Travesa et al., 2008). Another possibility to explain the lack of an evident cell cycle re-entry defect in the absence of PP4 during the induction of a reparable DSB is that other phosphatases might compensate the loss of PP4 during recovery. In this regard, it has been proposed that only the combination of *pph3Δ* with *ptc2*Δ and *ptc3*Δ restrains cells to deactivate Rad53 and resume the cell cycle (Travesa et al., 2008).

Apart of the role of PP4 in cell cycle resumption upon DNA repair, an intriguing question is how Rad53 hyper-phosphorylation in the absence of *PPH3* influences the repair process. Our data show that PP4 is required for an optimal DNA resection by reducing Rad53 phosphorylation levels, a mechanism that has been previously demonstrated to negative influence the activity of the pathway. Importantly, a double mutant *pph3Δ exo1*Δ was significantly more affected in resection than single mutants *pph3Δ* or *exo1*Δ, suggesting that PP4 contribution over Exo1 is not the main mechanism controlling resection. Still, we observed increased levels of Exo1 phosphorylation in the absence of PP4 activity during the DNA damage response implying that, even though PP4 can overcome Exo1 phosphorylation, this effect does not constitute a robust mechanism to inhibit its activity. In agreement with this observation, it has been reported that changing Exo1 phospho-residues to alanines or glutamic acids, to prevent or mimic phosphorylation respectively, has a subtle effect on Exo1 activity (Morin et al., 2008). Besides, the elevated phosphorylation observed in Exo1 in the absence of PP4 activity was completely abolished when introducing a Rad53 kinase dead allele, suggesting that PP4 counteracts Rad53 activity to restrain Exo1 phosphorylation. However, we cannot discard a direct function of PP4 in dephosphorylating directly Exo1 during the DDR to restrain its activity during the repair of a DNA lesion. Interestingly, combination of *pph3Δ* with *sgs1*Δ only slightly exacerbated the resection problems observed in the single mutants, suggesting that PP4-dependent Rad53 dephosphorylation predominantly affects Sgs1/Dna2 activity. It has been previously described that Rad9 provides a barrier to resection through limiting the association of Sgs1 to the DSB ends (Bonetti et al., 2015; Ferrari et al., 2015). Curiously, Rad9 association to the DSB vicinity depends on Rad53 kinase activity (Gobbini et al., 2015). Taking into account that we observed an increased Rad53-dependent phosphorylation of Rad9 in the absence of PP4 activity, we can conclude that Rad53 dephosphorylation by PP4 enhances Sgs1/Dna2-dependent resection by reducing Rad9 phosphorylation. Nevertheless, we cannot discard that PP4 might also be directly targeting Rad9 in order to reduce its negative effect on resection during the DNA damage response.

An important question that arises from these observations is how does the PP4/Rad53 module operate in the regulation of Rad9 association to damaged DNA. It has been previously reported that Rad9 binding to a DNA lesion depends specifically on H2A phosphorylation at S129 (Hammet et al., 2007). Chromatin-bound Rad9 is phosphorylated by Mec1 (Schwartz et al., 2002) to activate the recruitment of Rad53 to the DSB, an interaction that triggers its auto-phosphorylation activity (Pellicioli and Foiani, 2005). It has been reported in both human and yeasts that Rad9 binds to a DNA damaged site throughout its TUDOR domain (Grenon et al., 2007; Huyen et al., 2004). Taking into account that Rad9 hyper-phosphorylation in the absence of PP4 was mainly accumulated around the TUDOR domain, it is tempting to propose that Rad53-dependent phosphorylation of this region enhances Rad9 binding to chromatin. In this regard, PP4 activity over Rad53 could relax the interaction of Rad9 to damaged DNA in order to overcome the negative effect that this factor exerts over Sgs1/Dna2. This model fits perfectly well with the observation that elimination of S129 from H2A is sufficient to rescue both the DNA resection and repair phenotypes of cells lacking PP4 activity.

Even though Rad53-dependent Rad9 phosphorylation could constitute a perfect mechanism to explain the link between PP4 and resection we cannot discard that other targets of the phosphatase could be regulating the binding of Rad9 to the DNA lesion. In fact, it has been reported that the direct dephosphorylation of H2A by PP4 regulates not only cell cycle recovery (Keogh et al., 2006; Nakada et al., 2008) but also DNA repair (Chowdhury et al., 2008; Kim et al., 2011). Curiously, we found that H2A phosphorylation was indeed increased in the absence of PP4 activity in our MS screening during the repair of a DNA lesion. This result corroborates previous observations demonstrating that H2A is hyper-phosphorylated in cells lacking PP4 activity in response to MMS treatment (Kim et al., 2011). Taking into account that H2A phosphorylation is a prerequisite to recruit Rad9 to a DSB, it is feasible to think that the direct dephosphorylation of H2A by PP4 could be part of an additional mechanism that regulates Rad9 interaction to the DNA lesion. It is important to remark that Sgs1 phosphorylation by Mec1 has been implicated in the recruitment of Rad53 to damaged sites (Hegnauer et al., 2012). Therefore, we cannot discard that PP4 could also have a role in Rad9 regulation by acting over this helicase. Accordingly, we have also detected high levels of Sgs1 phosphorylation in the absence of PP4 activity in our mass spectrometry analysis. Whether the direct regulation of these targets by PP4 constitutes additional mechanisms to modulate the steady-state activity of the resection process along the damage response is a fascinating question for the future.

In addition to the molecular details behind PP4 regulation during the damage response, it is also important to focus our attention into the physiological significance of this phosphatase in controlling DNA end resection and its implications in DNA repair. Our data using an inter-chromosomal DSB repair approach mediated by gene conversion demonstrate that lack of PP4 activity does not influence DNA repair by ectopic recombination. This result confirms previous observations indicating that only the simultaneous depletion of PP4 and PP2C activities affects DNA repair through DSBR (Kim et al., 2011). As HR by gene conversion has been demonstrated to require limited amount of ssDNA (Cejka et al., 2010; Jinks-Robertson et al., 1993; Mimitou and Symington, 2008; Zhu et al., 2008), it is possible that PP4-dependent resection might be crucial for the repair of DSBs that rely on long-range resection. Accordingly, we have shown that PP4 activity becomes essential when cells are forced to use SSA as the unique pathway to restore a DSB. What is the benefit to support SSA by promoting long-range resection? Resection is activated in S phase through the increasing activity of the CDK, a mechanism that bias DNA repair towards HR even in the absence of the replicated chromatid. Thus, a DSB that occurs in S phase prior sister chromatid synthesis relies exclusively on SSA for its repair (Bhargava et al., 2016). In this context, repair by SSA might be critical for cell viability in response to DNA damage. It is important to remark that SSA is especially important in higher eukaryotes, whose genomes are enriched in DNA repeats. In this regard, it has been speculated that an inefficient resection in the absence of PP4 could lead to an excessive initiation of homologous recombination pathways that increase the accumulation of DNA intermediates and rearrangements between repeats, a feature that has been proven to increase genome heterozygosis (Weinstock et al., 2006). Additionally, it has been demonstrated that reliance on SSA could compensate genetic deficiencies in HR pathways, as the BRCA2 mutation associated with breast/ovarian cancer (Tutt et al., 2001).

In conclusion, we propose a model whereby the phosphorylation of Rad53 is tightly controlled by its own kinase activity and the PP4 phosphatase during the DNA damage response. The role of PP4 in counteracting Rad53 auto-activation is essential to restrain the inhibitory role that Rad9 exerts over the Sgs1/Dna2 pathway, thus triggering a robust and efficient resection process. This function becomes vital when repairing a DNA lesion by mechanisms that rely on extensive resection for its achievement. Importantly, this model implies that excessive checkpoint activation is not compatible with DNA repair pathways that depend on long-range resection for their accomplishment and envisions that DDR kinases and phosphatases must cooperate along the damage response to couple checkpoint activation with DNA repair. This balance in checkpoint activation ensures a precise DNA damage response, strong enough to activate an accurate G2/M cell cycle arrest but not too robust to negatively influence in the repair of a DNA lesion.

## Experimental Procedures

### Yeast strains, growing conditions and plasmids

All strains used in this study are listed in Appendix Table S1. Epitope tagging of endogenous genes was performed by gene targeting using polymerase chain reaction (PCR) products as described in (Janke et al., 2004). For the majority of the experiments cells were grown in YEP containing 2% raffinose. HO expression was attained by adding galactose to a final concentration of 2%. Samples were collected for FACS, DNA and protein analysis before adding galactose and at different time points after the HO induction. The *rad53K227A* mutation was introduced using an *Eco*RI-linearized pCH3 plasmid (Pellicioli et al., 1999). All strains containing the *hta1/hta2-S129\** versions of H2A were constructed as previously described (Downs et al., 2000).

### Southern blots

Cell lysis was achieved by treating the samples with 40 units of lyticase in DNA preparation buffer (1% SDS, 100 μM NaCl, 50 μM Tris-HCl, 10 μM EDTA) for 10 min. DNA extraction was performed by mixing the samples with phenol:chloroform:isoamylalcohol (25:24:1) for 10 min. After centrifugation, the soluble fraction was precipitated with ethanol and resuspended in TE buffer. Purified DNA was digested with the appropriate restriction enzyme, separated on 0.7%-1% agarose gels and subjected to Southern blotting. Probing was performed by labelling a PCR-amplified DNA fragment with a mix of nucleotides containing Fluorescein-12-dUTP (Fluorescein-High Prime, Roche). The oligonucleotides used to synthesise the probes are included in Appendix Table S2. Detection was attained by using an anti-Fluorescein antibody conjugated with alkaline phosphatase (Anti-Fluorescein-AP Fab fragments, Roche). A FIJI software was used for the processing, analysis and quantification of the images.

### Western Blots

Samples were prepared by using a trichloroacetic acid (TCA) method. In brief, cells were collected by centrifugation and fixed with 20% TCA. Cells were broken by shaking the pellet with glass beads in a FastPrep machine (MPBio) (3×20 sec cycles at power setting 5.5). Protein precipitation was reached by centrifugation at 5,000 rpm for 5 min at 4ºC. Upon precipitation, the pellet was resuspended in 1M Tris-HCl pH 8 and SDS-PAGE loading buffer, and boiled for 10 min. Insoluble material was separated by centrifugation and the supernatant was loaded in a 6% acrylamide gel. When separating Rad9 phospho-bands, gels were supplemented with 5 μM of Phos-Tag and 10 μM of MnCl_2_. For Exo1 phosphorylation analysis, 20 μM of Phos-Tag and 40 μM of MnCl_2_ were used. Proteins were transferred into PVDF membranes (Hybond-P, GE Healthcare) and blocked with 5% milk in PBS-Tween (0.1%). Anti-Rad53 antibody was used at a 1:2,000 dilution (Abcam) and the secondary anti-rabbit antibody was used at a concentration of 1:25,000 (GE Healthcare). Both anti-HA and anti-MYC antibodies were used at a 1:2,500 dilution (Roche), and the secondary anti-mouse antibody was used at a concentration of 1:25,000 (GE Healthcare). After several washes in PBS-Tween, membranes were incubated with SuperSignal^®^ West Femto (Thermo Scientific), followed by exposure to ECL Hyperfilm (GE Healthcare).

### Liquid chromatography-tandem mass spectrometry (LC-MS/MS) analysis

For each sample, 150 ODs of cells were harvested at 4°C, washed once in cold water and frozen at -80°C. Cell pellets were thawed briefly on ice and resuspended in 150 µl of 8 M Urea, 20 mM HEPES pH 8 buffer supplemented with PhosSTOP phosphatase inhibitor tablets (Roche). 300 µl of 425-600 µm glass beads (Sigma) were added and protein extracts were prepared by bead beating using a FastPrep machine (MPBio) at 3×30 sec pulses and 5.5 power setting. Cell lysates were resuspended in 300 µl of fresh buffer and the soluble fraction was separated from the insoluble fraction by centrifugation at 15,000 rpm for 15 min at 4°C.

For sample processing, 550 µg of protein were reduced and alkylated sequentially with 10 mM dithiothreitol and 50 mM 2-chloroacetamide respectively. Samples were then diluted with 20 mM HEPES buffer (pH 8.0) to 4 M urea and sufficient LysC protease (Wako, 125-05061) was added for a protease to protein ratio of 1:250, followed by incubation (37°C, 5 hours). Samples were then diluted to 1 M urea and trypsin (Promega, V5280) was added to a 1:50 ratio for overnight incubation at 37°C. Samples were acidified with trifluoroacetic acid (TFA) to a final concentration of 0.5% prior to solid phase extraction. For total protein analysis, volumes equivalent to 50 µg of protein were desalted using C18 spin tips (Glygen Corp, TT2C18.96) and peptides eluted with 60% ACN + 0.1% formic acid (FA). Eluents were then dried using a centrifugal vacuum drier. The remaining sample volumes were de-salted using solid phase extraction with OASIS HLB 10 mg cartridges (Waters, 186000383) according to the manufacturer’s instructions.

Peptides were eluted from OASIS HLB cartridges with 1 M glycolic acid in 80% acetonitrile (ACN), 5% TFA and phospho-enriched using a TiO_2_ based method (Casado et al., 2014). Briefly, eluents were adjusted to 1 mL with 1 M glycolic acid solution and were then incubated with 25 mg of TiO_2_ (50% slurry in 1% TFA) for 5 min at room temperature. After 5 min of incubation with mixing, the TiO_2_/peptide mixture was packed into empty spin tips by centrifugation. The TiO_2_ layer was then sequentially washed with 1 M glycolic acid solution, 100 mM ammonium acetate in 25% ACN and 10% ACN. Phosphopeptides were eluted with 4 sequential additions of a 5% NH_4_OH solution. Eluents were clarified with centrifugation at 13,000 rpm for 2 min and clear supernatant transferred to new tubes. Samples were snap frozen on dry ice and then dried by vacuum centrifugation.

Phospho-enriched samples and desalted total protein digests were redissolved in 0.1% TFA by shaking (1,200 rpm) for 30 min and sonication on an ultrasonic water bath for 10 min, followed by centrifugation (14,000 rpm, 5°C) for 10 min. LC-MS/MS analysis was carried out in technical duplicates (1 µg on column for total protein samples) and separation was performed using an Ultimate 3000 RSLC nano liquid chromatography system (Thermo Scientific) coupled to an Orbitrap Velos mass spectrometer (Thermo Scientific) via a Dionex nano-electrospray source. For LC-MS/MS analysis, phosphopeptide solutions were injected and loaded onto a trap column (Acclaim PepMap 100 C18, 100 µm × 2 cm) for desalting and concentration at 8 µL/min in 2% acetonitrile, 0.1% TFA. Peptides were then eluted on-line to an analytical column (Acclaim Pepmap RSLC C18, 75 µm × 50 cm) at a flow rate of 250 nL/min. Peptides were separated using a 120 minute gradient, 4-25% of buffer B for 90 min followed by 25-45% buffer B for another 30 min (buffer A: 5% DMSO, 0.1% FA; buffer B: 75% acetonitrile, 5% DMSO, 0.1%FA) and subsequent column conditioning and equilibration. Eluted peptides were analysed by the mass spectrometer operating in positive polarity using a data-dependent acquisition mode. Ions for fragmentation were determined from an initial MS1 survey scan at 30,000 resolution, followed by CID (Collision-Induced Dissociation) of the top 10 most abundant ions. MS1 and MS2 scan AGC targets were set to 1^6^ and 3^4^ for maximum injection times of 500 ms and 100 ms respectively. A survey scan m/z range of 350-1500 was used, with multistage activation (MSA) enabled, normalised collision energy set to 35%, charge state screening enabled with +1 charge states rejected and minimal fragmentation trigger signal threshold of 500 counts.

The data was processed using the MaxQuant software platform (v1.6.1.0), with database searches carried out by the in-built Andromeda search engine against the Uniprot *S. cerevisiae* database (version 20180305, number of entries: 6,729). A reverse decoy database approach was used at a 1% false discovery rate (FDR) for peptide spectrum matches. Search parameters included: maximum missed cleavages set to 2, fixed modification of cysteine carbamidomethylation and variable modifications of methionine oxidation, protein N-terminal acetylation and serine, threonine, tyrosine phosphorylation. For total protein samples, search parameters as above with variable modifications of methionine oxidation, protein N-terminal acetylation, asparagine deamidation and cyclization of N-terminal glutamine to pyroglutamate. Label-free quantification was enabled with an LFQ minimum ratio count of 2. ‘Match between runs’ function was used with match and alignment time limits of 1 and 20 min respectively.

## Statistical analysis

All quantifications shown in the article represent the mean ± SD from at least three independent experiments. The data were statistically analysed using a *t-*test and the differences between the strains tested were indicated in the graphs unless otherwise stated (*P*<0.05*, *P*<0.005**, *P*<0.0005***). GraphPad Prism software was used for all statistical analysis.

## Author Contribution

A.C-B. planned and evaluated most experiment. M.T.V. was responsible for the development of the majority of the experiments reported in the paper. P.G.E. and L.A. were responsible for the mass spectrometry experiments. A.M. and H.K. were responsible for performing the mass spectrometry analysis. F.R. and E.M. provided technical support to this work. A.C-B. wrote the paper and all authors analysed the data, discussed the results and commented on the manuscript.

## Acknowledgments

We would like to thank P. San Segundo and J. Haber for providing valuable strains and plasmids. We particularly thank Vincent Géli and Jessica A. Downs for providing plasmids. We thank P. San Segundo, CR. Vazquez de Aldana and J. Correa for helpful comments, discussions and reading of the manuscript. This work was funded by grants from the “Ministerio de Economía y Competitividad” (BFU2013-41216-P and BFU2016-77081-P) conceded to A.C-B. The IBFG is supported in part by an institutional grant from the “Junta y Castilla y León” (CLU-2017-03). M.T.V. was recipient of a predoctoral fellowship from the “Junta de Castilla y León”. F.R. was recipient of a predoctoral fellowship from the “Ministerio de Economía y Competitividad”.

## Appendix Tables

**Appendix Table S1.** Genotypes of strains used in this study.

| Strain | Genotype | Reference |
| --- | --- | --- |
| 218 | HOΔ *hml::ADE1 mata::hisG hmr::ADE1 his4::NAT-leu-(Xho- to Asp718) leu2::HOcs ade3::GAL::HO ade1 lys5 ura3-52 trp1::hisG*. | J. Haber |
| 224 | MATa-inc HOΔ *hml::ADE1 hmr::ADE1 ade1-100 leu2-3,112 lys5 trp1::hisG ura3-52 ade3::GAL::HO arg5,6::MATa::HPH* | J. Haber |
| 406 | HOΔ MATa *ade1-100 leu2,3-112 lys5 ura3-52 trp1::hisG hml::ADE1 hmr::ADE1 ade3::GAL-HO* | J. Haber |
| 825 | HOΔ MATa *ade1-100 leu2,3-112 lys5 ura3-52 trp1::hisG hml::ADE1 hmr::ADE1 ade3::GAL-HO exo1Δ::TRP1* | L. Aragón |
| 1327 | HOΔ *hml::ADE1 mata::hisG hmr::ADE1 his4::NAT-leu-(Xho- to Asp718) leu2::HOcs ade3::GAL::HO ade1 lys5 ura3-52 trp1::hisG pph3Δ::HPH* | This Study |
| 1344 | HOΔ *hml::ADE1 MATa-inc hmr::ADE1 ade1 leu2-3,112 lys5 trp1::hisG ura3-52 ade3::GAL::HO arg5,6::MATa::HPH pph3Δ::LEU* | This Study |
| 1366 | HOΔ MATa *ade1-100 leu2,3-112 lys5 ura3-52 trp1::hisG hml::ADE1 hmr::ADE1 ade3::GAL-HO pph3Δ::HPH* | This Study |
| 1420 | HOΔ MATa *ade1-100 leu2,3-112 lys5 ura3-52 trp1::hisG hml::ADE1 hmr::ADE1 ade3::GAL-HO exo1Δ::TRP1 pph3Δ::HPH* | This Study |
| 1468 | HOΔ *hml::ADE1 mata::hisG hmr::ADE1 his4::NAT-leu-(Xho- to Asp718) leu2::HOcs ade3::GAL::HO ade1 lys5 ura3-52 trp1::hisG rad51Δ::URA* | This Study |
| 1471 | HOΔ *hml::ADE1 mata::hisG hmr::ADE1 his4::NAT-leu-(Xho- to Asp718) leu2::HOcs ade3::GAL::HO ade1 lys5 ura3-52 trp1::hisG pph3Δ::HPH rad51Δ::URA* | This Study |
| 1473 | HOΔ *hml::ADE1 mata::hisG hmr::ADE1 his4::NAT-leu-(Xho- to Asp718) leu2::HOcs ade3::GAL::HO ade1 lys5 ura3-52 trp1::hisG rad53-K227A::KanMX* | This Study |
| 1476 | HOΔ *hml::ADE1 mata::hisG hmr::ADE1 his4::NAT-leu-(Xho- to Asp718) leu2::HOcs ade3::GAL::HO ade1 lys5 ura3-52 trp1::hisG pph3Δ::HPH rad53-K227A::KanMX* | This Study |
| 1494 | HOΔ MATa *ade1-100 leu2,3-112 lys5 ura3-52 trp1::hisG hml::ADE1 hmr::ADE1 ade3::GAL-HO pph3Δ::HPH rad53-K227A::KanMX* | This Study |
| 1496 | HOΔ *hml::ADE1 mata::hisG hmr::ADE1 his4::NAT-leu-(Xho- to Asp718) leu2::HOcs ade3::GAL::HO ade1 lys5 ura3-52 trp1::hisG rad51Δ::URA rad53-K227A::KanMX* | This Study |
| 1502 | HOΔ *hml::ADE1 mata::hisG hmr::ADE1 his4::NAT-leu-(Xho- to Asp718) leu2::HOcs ade3::GAL::HO ade1 lys5 ura3-52 trp1::hisG pph3Δ::HPH rad51Δ::URA rad53-K227A::KanMX* | This Study |
| 1536 | HOΔ *hml::ADE1 mata::hisG hmr::ADE1 his4::NAT-leu-(Xho- to Asp718) leu2::HOcs ade3::GAL::HO ade1 lys5 ura3-52 trp1::hisG rad9Δ::KanMX* | This Study |
| 1538 | HOΔ *hml::ADE1 mata::hisG hmr::ADE1 his4::NAT-leu-(Xho- to Asp718) leu2::HOcs ade3::GAL::HO ade1 lys5 ura3-52 trp1::hisG pph3Δ::HPH rad9Δ::KanMX* | This Study |
| 1541 | HOΔ *hml::ADE1 mata::hisG hmr::ADE1 his4::NAT-leu-(Xho- to Asp718) leu2::HOcs ade3::GAL::HO ade1 lys5 ura3-52 trp1::hisG rad51Δ::URA rad9Δ::KanMX* | This Study |
| 1543 | HOΔ *hml::ADE1 mata::hisG hmr::ADE1 his4::NAT-leu-(Xho- to Asp718) leu2::HOcs ade3::GAL::HO ade1 lys5 ura3-52 trp1::hisG pph3Δ::HPH rad51Δ::URA rad9Δ::KanMX* | This Study |
| 1547 | HOΔ MATa *ade1-100 leu2,3-112 lys5 ura3-52 trp1::hisG hml::ADE1 hmr::ADE1 ade3::GAL-HO pph3Δ::HPH rad9Δ::KanMX* | This Study |
| 1557 | HOΔ MATa *ade1-100 leu2,3-112 lys5 ura3-52 trp1::hisG hml::ADE1 hmr::ADE1 ade3::GAL-HO Rad9-6HA::KanMX* | This Study |
| 1559 | HOΔ MATa *ade1-100 leu2,3-112 lys5 ura3-52 trp1::hisG hml::ADE1 hmr::ADE1 ade3::GAL-HO* *pph3Δ::HPH Rad9-6HA::KanMX* | This Study |
| 1660 | HOΔ MATa *ade1-100 leu2,3-112 lys5 ura3-52 trp1::hisG hml::ADE1 hmr::ADE1 ade3::GAL-HO* *Exo1-9myc::NAT* | This Study |
| 1661 | HOΔ MATa *ade1-100 leu2,3-112 lys5 ura3-52 trp1::hisG hml::ADE1 hmr::ADE1 ade3::GAL-HO* *pph3Δ::HPH Exo1-9myc::NAT* | This Study |
| 1665 | HOΔ MATa *ade1-100 leu2,3-112 lys5 ura3-52 trp1::hisG hml::ADE1 hmr::ADE1 ade3::GAL-HO* *pph3Δ::HPH rad53-K227A::KanMX Exo1-9myc::NAT* | This Study |
| 1773 | HOΔ MATa *ade1-100 leu2,3-112 lys5 ura3-52 trp1::hisG hml::ADE1 hmr::ADE1 ade3::GAL-HO* *pph3Δ::HPH rad53-K227A::KanMX Rad9-6HA::NAT* | This Study |
| 1792 | HOΔ MATa *ade1-100 leu2,3-112 lys5 ura3-52 trp1::hisG hml::ADE1 hmr::ADE1 ade3::GAL-HO* *sgs1Δ::NAT* | This Study |
| 1794 | HOΔ MATa *ade1-100 leu2,3-112 lys5 ura3-52 trp1::hisG hml::ADE1 hmr::ADE1 ade3::GAL-HO* *pph3Δ::HPH sgs1Δ::NAT* | This Study |
| 1883 | HOΔ *hml::ADE1 mata::hisG hmr::ADE1 his4::NAT-leu(Xho- to Asp718) leu2::HOcs ade3::GAL::HO* *ade1 lys5 ura3-52 trp1::hisG pph3Δ::HPH hta1-S129\*::hisG hta2-S129\*::hisG* | This Study |

**Appendix Table S2.**
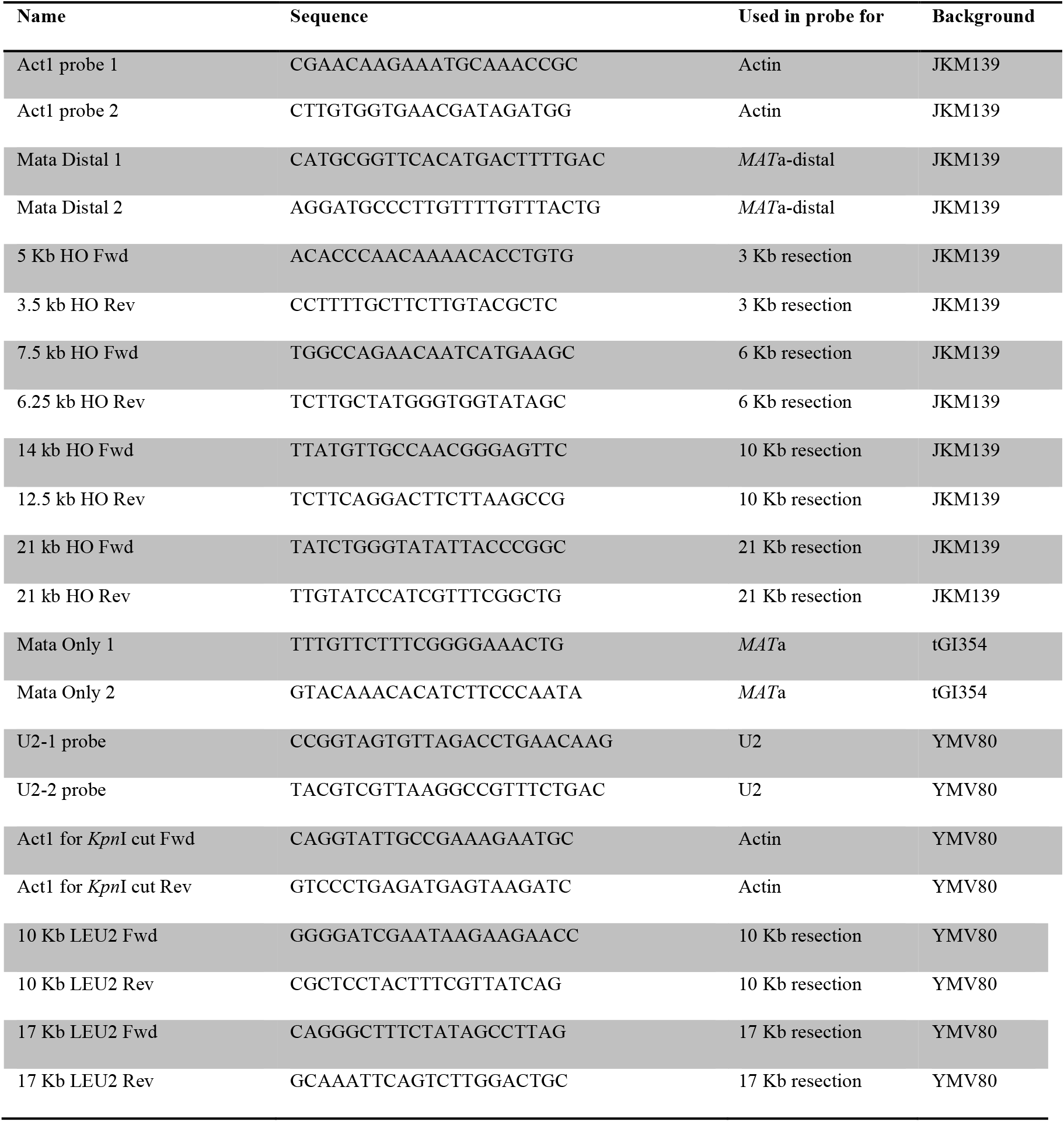
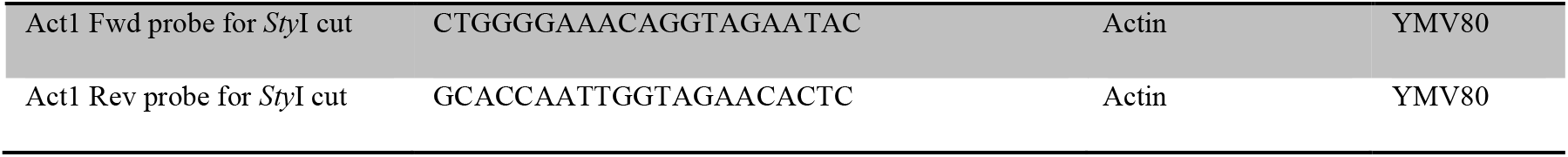
Oligonucleotides used for probes synthesis.

**Appendix Table S3.**
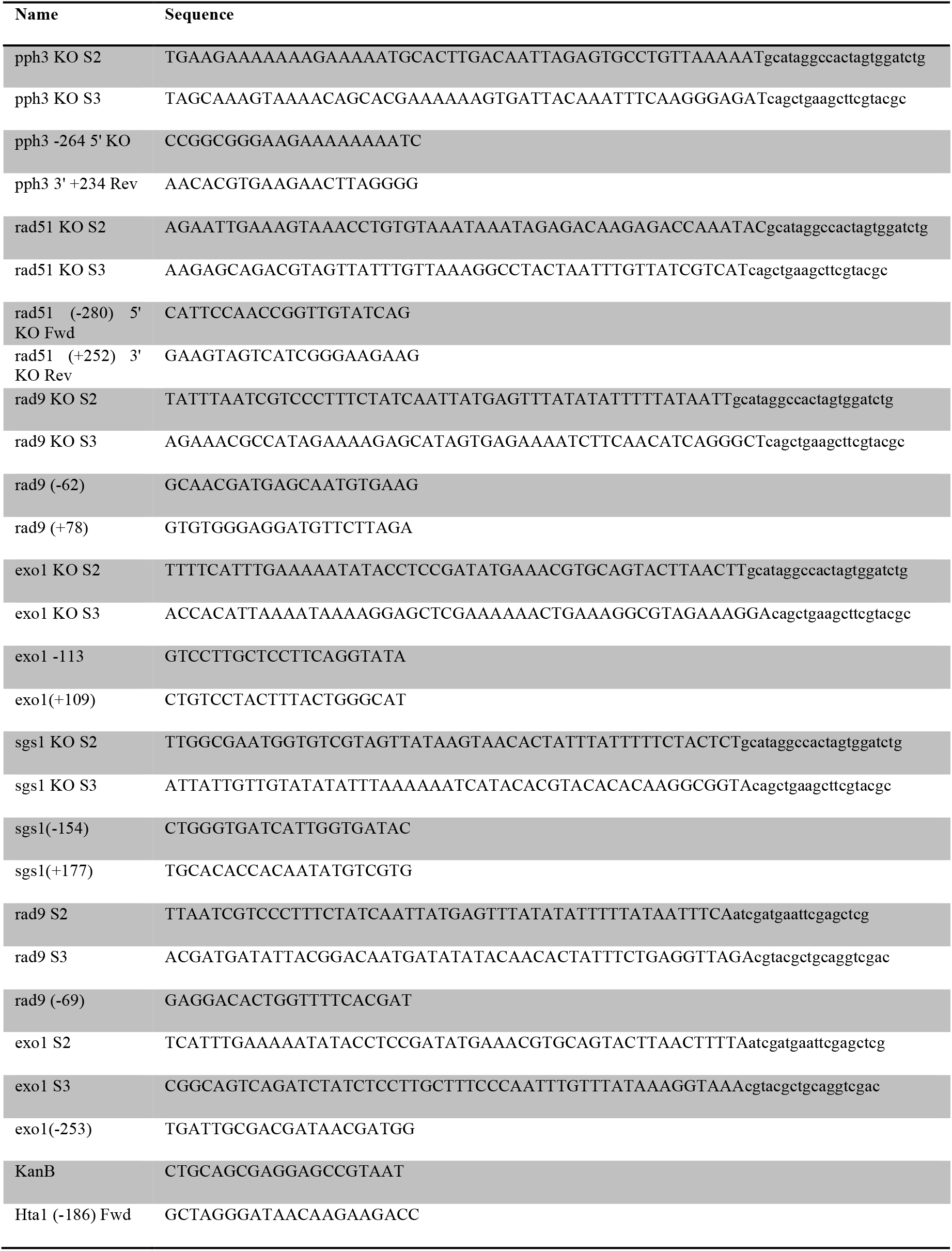
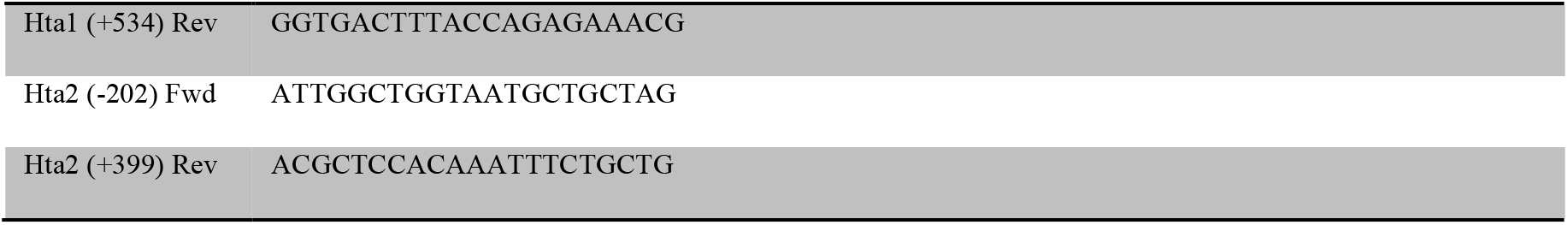
Oligonucleotides used for strain generation.

---

**Supplementary Figure 1.**
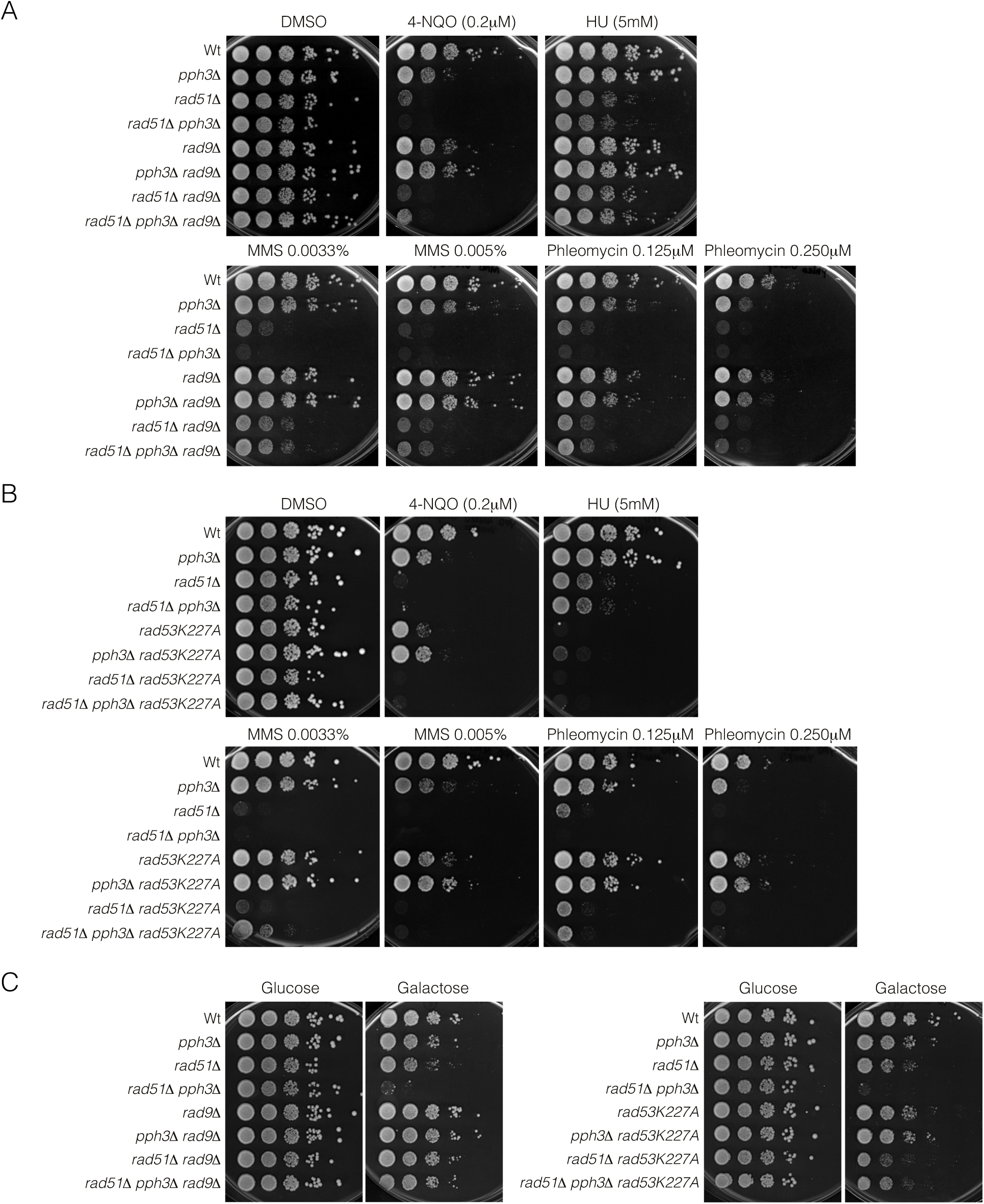
Drops assays to evaluate the sensitivity of different genetic backgrounds to several sources of DNA damage. A) Deletion of Rad9 improves cell viability of cells lacking PP4 activity. Ten-fold serial dilutions from overnight cultures of the indicated strains dropped and grown on solid rich media containing mock DMSO (as non-treated control), 4-NQO (0.2 μM), HU (5 μM), MMS (0.0033% and 0.005%) and Phleomycin (0.125 μM and 0.250 μM). B) Elimination of Rad53 kinase activity increases cell viability of PP4-deficient cells. Ten-fold serial dilutions from mid-log phase cultures of the indicated strains grown as in A). C) Disruption of Rad9 or Rad53 kinase activity restores cell viability of *pph3Δ* in response to a single DSB. Exponential overnight cell cultures of the specified strains were serial diluted (1:10) and each dilution was plated on 2% glucose (no DSB) or galactose (DSB).

**Supplementary Figure 2.**
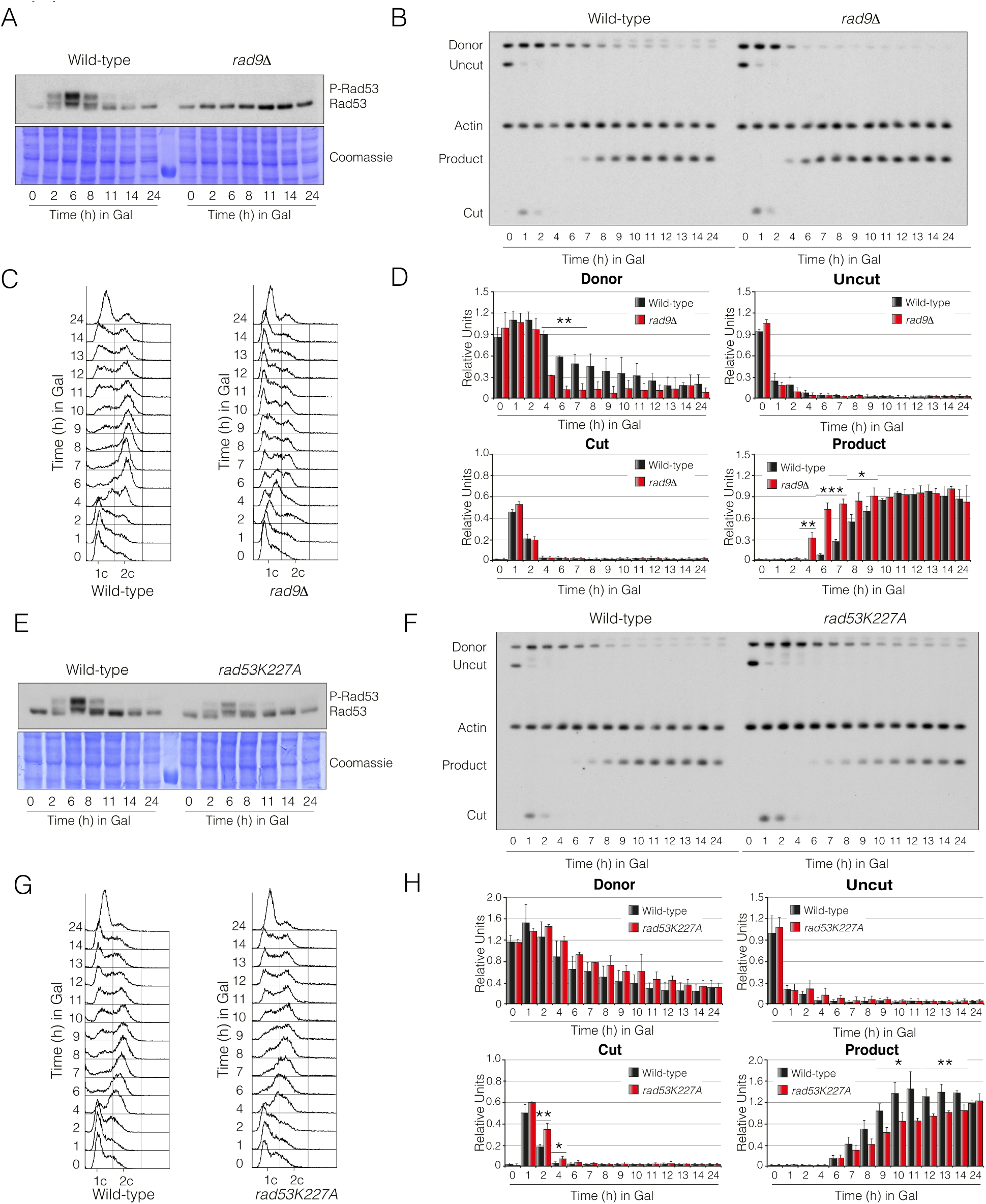
Lack of Rad9 or Rad53 activities differentially affects DNA repair by SSA/BIR. A) Elimination of Rad9 drastically affects Rad53 phosphorylation. Exponentially wild-type and *rad9*Δ cells growing in YP-Raffinose were HO-induced by adding galactose to the media and samples were taken at the indicated time points. Proteins were TCA extracted and subjected to Western blotting. Coomassie staining is shown as loading control. B) Lack of Rad9 slightly enhances DNA repair by SSA/BIR. Southern blot analysis of wild-type and *rad9*Δ cells harbouring an YMV80-based repair system. Mid-log cells growing in YP-Raffinose were supplemented with galactose and samples were taken at the specified time points. DNA was extracted, digested with *Kpn*I and blotted. Blots were probed with an *U2* DNA sequence and *ACT1* as loading control. C) FACS profile for DNA content of samples taken from the experiment shown in B). D) Graphs represent the quantification of the band signals of the Southern blot depicted in B). All data were normalized using the actin signal as loading control. Graphs represent the mean ± SD from three independent experiments. P-values were calculated using a two-tailed unpaired Student’s t-test. E) Depletion of Rad53 auto-phosphorylation reduces its phosphorylation state along the damage response. A single DSB was induced in mid-log wild-type and *rad53K227A* cells growing in YP-Raffinose by adding galactose to the media and samples were taken at the indicated time points. Proteins were TCA extracted and subjected to Western blotting. Coomassie staining is shown as loading control. F) Rad53 auto-phosphorylation is needed for DNA repair by SSA/BIR. Wild-type and *rad53K227A* strains containing an YMV80-based repair system were grown in YP-Raffinose and supplemented with galactose. Samples at the indicated time points were taken and subjected to Southern blot analysis. Genomic DNA was digested with *Kpn*I and blotted. Blots were probed with an *U2* DNA sequence and *ACT1* as loading control. G) FACS profile for DNA content of samples taken from the experiment shown in F). H) Graphs represent the quantification of the band signals detected in the Southern blot depicted in F). All data were normalized using the actin signal as loading control. Graphs represent the mean ± SD from three independent experiments. P-values were calculated using a two-tailed unpaired Student’s t-test.

**Supplementary Figure 3.**
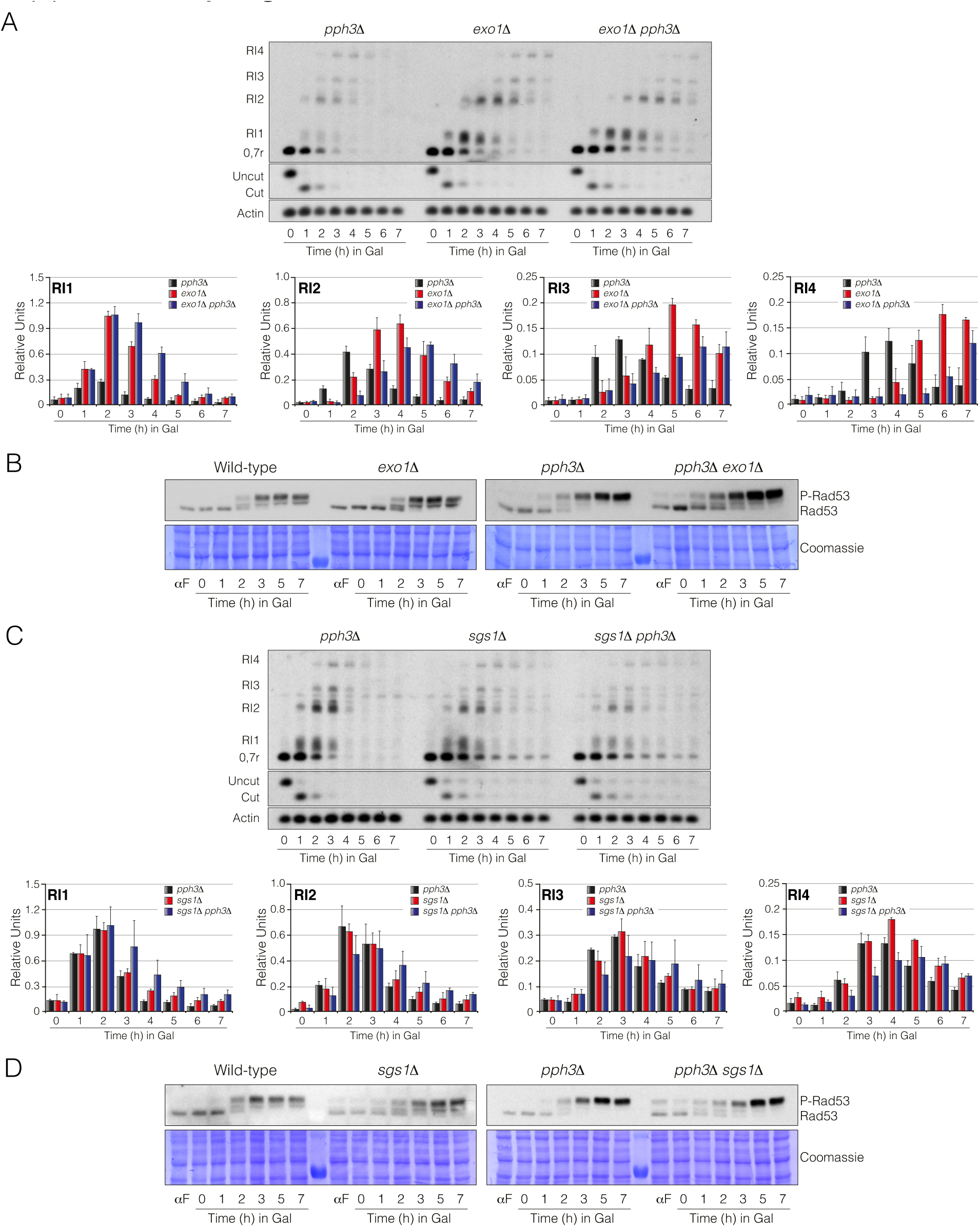
PP4 enhances DNA end resection by regulating the Sgs1/Dna2 pathway. A) Samples taken from experiment 5A) were DNA extracted, digested with *Sty*I and blotted. The density of the different resection intermediate signals was measured and plotted. All data were normalized using the actin signal as loading control. Graphs represent the mean ± SD from three independent experiments. B) Same experiment as in figure 5A) was TCA processed and subjected to Western blotting to detect the pattern of Rad53 phosphorylation along the induction of a non-reparable DSB in wild-type, *exo1*Δ, *pph3Δ* and *pph3Δ exo1*Δ background strains. C) Samples collected from experiment 5B) were DNA extracted, digested with *Sty*I and blotted. The intensity of the different resection intermediates bands was measured and plotted. All data were normalized using the actin signal as loading control. Graphs represent the mean ± SD from three independent experiments. D) Same experiment as in figure 5B) was TCA processed and subjected to Western blotting to determine the Rad53 phosphorylation profile along the induction of a non-reparable DSB in wild-type, *sgs1*Δ, *pph3Δ* and *pph3Δ sgs1*Δ strains.

**Supplementary Figure 4.**
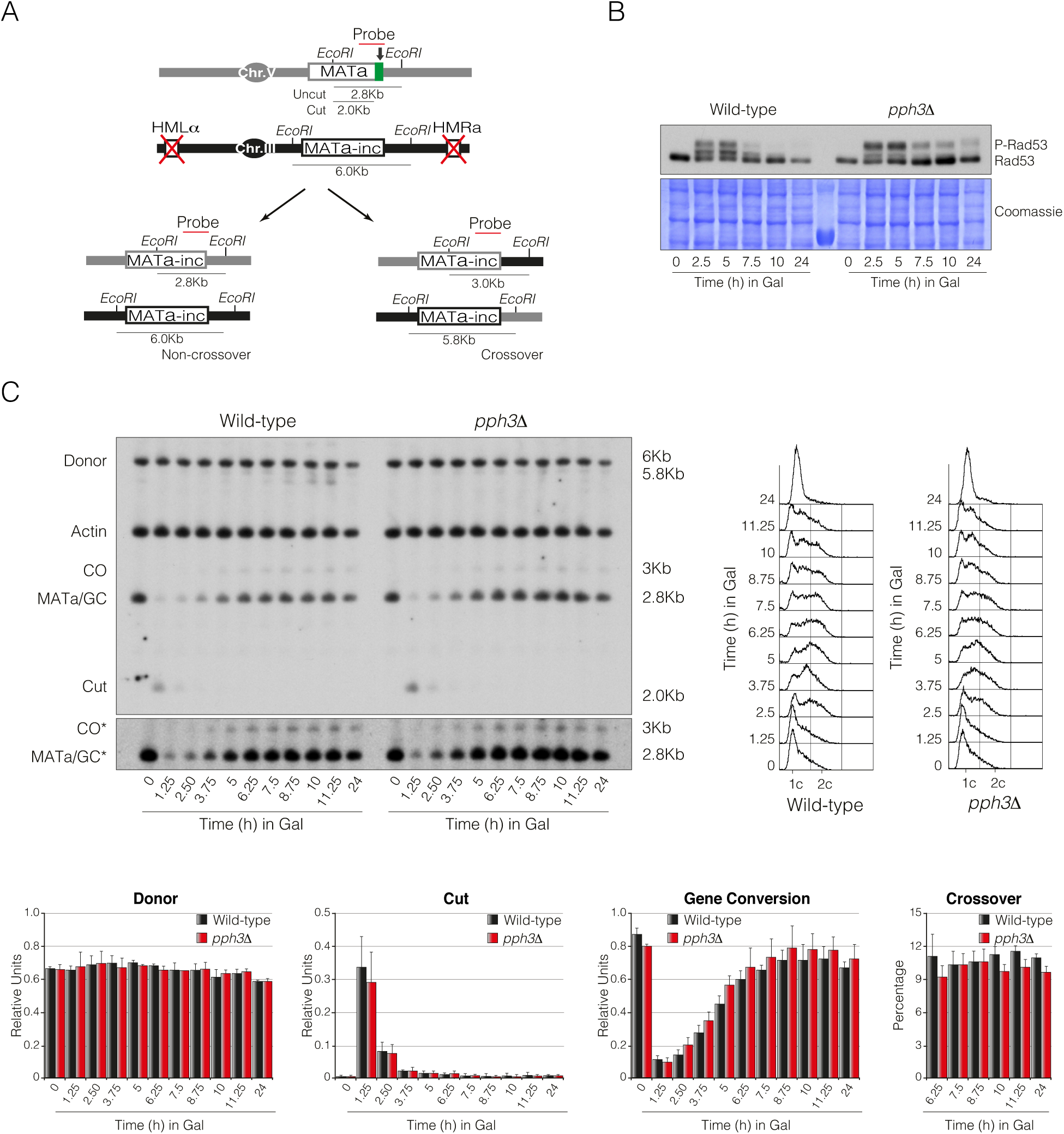
PP4 is not required for inter-chromosomal recombination. A) Schematic diagram displaying relevant genomic structure of the strain used to measure inter-chromosomal DNA repair proficiency. The localization of a *MAT-*specific probe and the restriction endonuclease cleavage sites used for Southern blot analysis to detect repair product formation are indicated. Note that crossover and non-crossover products have different restriction fragments that can be differentiated in Southern blot experiments. Arrow depicts the localization of the HO cleavage site. B) Immunoblot of Rad53 was performed in wild-type and *pph3Δ* cells to determine its phosphorylation level along the induction of the DSB. C) Physical analysis of wild-type and *pph3Δ* mutant strains carrying the inter-chromosomal gene conversion assay depicted in A). Cells were grown overnight before adding galactose to induce HO expression, thus producing a DSB on chromosome V. Samples were taken at different time points, and genomic DNA was extracted, digested with *Eco*RI and analysed by Southern blot. Blots were hybridized with a *MATa-*only DNA sequence, and an actin probe was used as loading control. Asterisk denotes an overexposed film to visualize crossover formation. FACS analyses are shown. Graphs represent the mean ± SD from three independent experiments. All data were normalized using actin as a loading control.

**Supplementary Figure 5.**
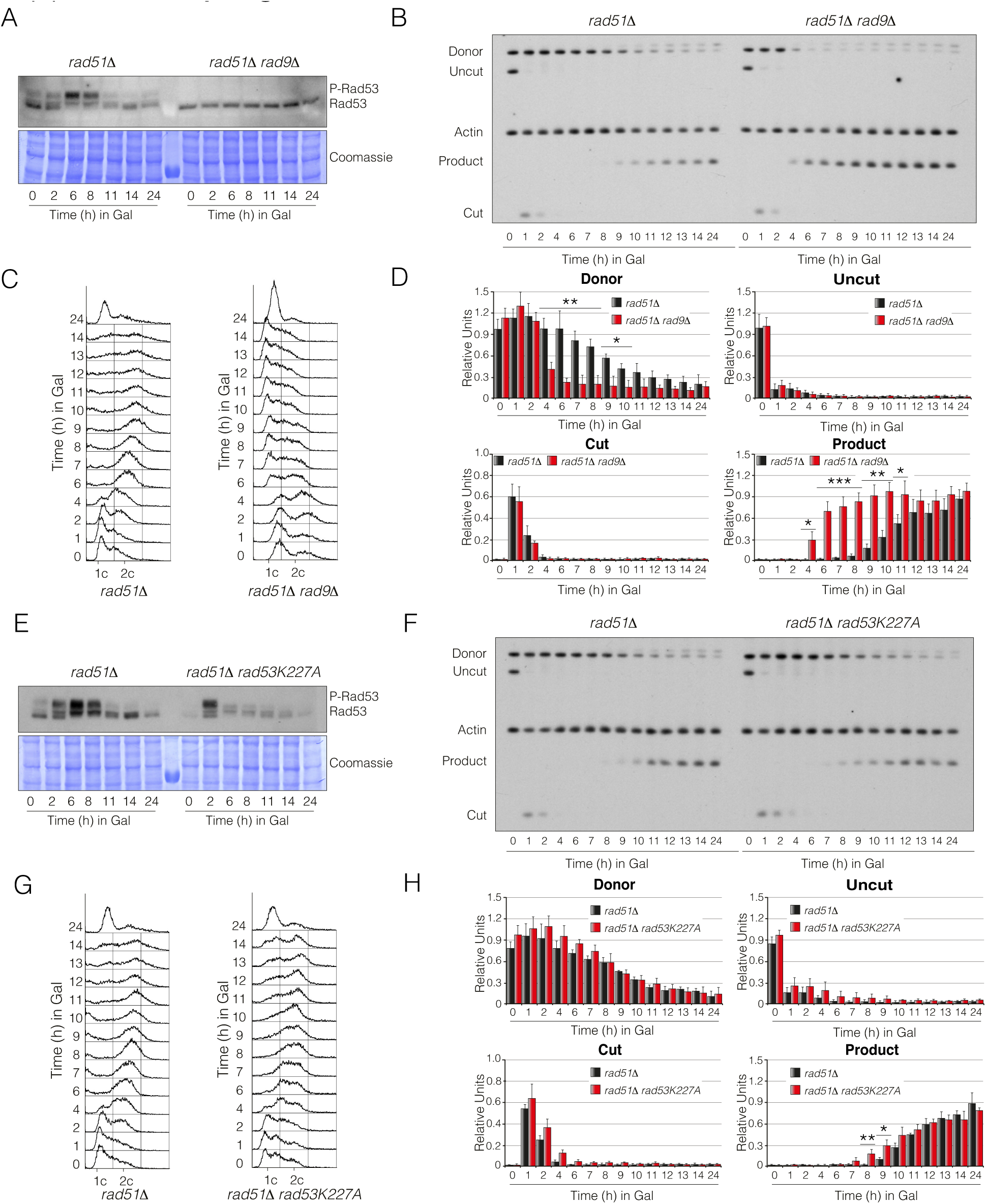
The steady-state levels of Rad53 phosphorylation are important for SSA execution. A) Disruption of Rad9 reduces Rad53 phosphorylation during SSA execution. Exponentially growing YP-Raffinose cell cultures of YMV80 derivative strains containing *rad51Δ* and *rad51Δ rad9*Δ deletions were supplemented with galactose to induce the expression of the HO endonuclease. Samples were collected at the indicated time points, TCA extracted and subjected to Western blotting. Coomassie staining is shown as loading control. B) Lack of Rad9 improves SSA repair efficiency. Physical analysis of *rad51Δ* and *rad51Δ rad9*Δ backgrounds containing the DNA repair assay portrayed in figure 7A. Cells were grown overnight in YP-Raffinose and supplemented with galactose. Samples were taken at different time points, genomic DNA extracted, digested with *Kpn*I and analysed by Southern blot. Blots were probed with an *U2* probe and *ACT1* was used as loading control. C) FACS profile for DNA content of samples collected from the experiment depicted in B). D) Graphs represent the quantification of the band signals obtained from the Southern blot shown in B). All experiments were normalized using the actin signal. Graphs represent the mean ± SD from three independent experiments. Replicates were averaged and statistical significance of differences assessed by a two-tailed unpaired Student’s t-test. E) Inactivation of the kinase activity of Rad53 affects its own phosphorylation during SSA execution. Cells from *rad51Δ* and *rad51Δ rad53K227A* background strains were grown overnight in raffinose-containing media and galactose was added to induce the expression of the HO endonuclease. Samples were collected at the indicated time points and subjected to Western blotting. Coomassie staining is shown as loading control. F) Elimination of Rad53 activity impairs DNA repair by SSA. Southern blot of *rad51Δ* and *rad51Δ rad53K227A* cells carrying the DNA repair system depicted in figure 7A. Exponentially growing YP-Raffinose cells were supplemented with galactose and samples were taken at the indicated time points. DNA was extracted, digested with *Kpn*I and blotted. Blots were probed with an *U2* DNA sequence and *ACT1* as loading control. G) FACS profile for DNA content of samples collected from experiment shown in F). H) Graphs representing the quantification of the band intensities obtained from the Southern blot depicted in F). All data were normalized using the actin signal. Graphs represent the mean ± SD from three independent experiments. P-values were calculated using a two-tailed unpaired Student’s t-test.

